# Exploring the relationship between vascular remodelling and tumour growth using agent-based modelling

**DOI:** 10.1101/2025.03.17.643670

**Authors:** Nicholas Fan, Joshua A. Bull, Helen M. Byrne

## Abstract

We develop a multiscale agent-based model (ABM) to investigate the effect that mechanical interactions between proliferating tumour cells and the surrounding vasculature have on the oxygen supply to the tumour microenvironment (TME), the tumour’s growth dynamics, and its response to radiotherapy. Our model extends existing models of tumour spheroid growth by incorporating vessel deformation due to mechanical forces between vessel walls and neighbouring tumour cells. These forces generate an effective pressure which compresses vessels, driving occlusion and pruning. This, in turn, leads to a hypoxic oxygen landscape which stimulates angiogenesis.

A key feature of our model is the treatment of mechanical cell interactions with the tumour microenvironment, which we represent with two forces. The first is Stokes’ drag which is widely used in ABMs to represent resistance to cell movement. The second is a friction force which accounts for resistance due to the continual breaking and reforming of cell-extracellular matrix (ECM) adhesions. The importance of this friction force is demonstrated by numerical simulation. When Stokes’ drag dominates, pressure gradients dissipate across the tissue and vessel compression is negligible. By contrast, as the strength of the friction force increases, larger pressure gradients form, leading to significant vessel compression.

We perform extensive numerical simulations to investigate how model parameters that control vascular remodelling and friction influence tumour vascularisation, which we spatially quantify using the cross-pair correlation function. This, in turn, alters the oxygen landscape and drives changes in tumour morphology. Finally, we highlight the importance of accounting for both mechanisms when simulating tumour responses to treatment with radiotherapy. We observe that vascular remodelling critically alters the tumour’s susceptibility to treatment and post-radiotherapy regrowth. Tumour regrowth is especially impacted by vessel remodelling, with certain vascular landscapes able to rebound quickly post-radiotherapy, resulting in fast tumour regrowth.

**Author Summary:** We have created an agent-based model (ABM) that accounts for mechanical interactions between tumour cells and associated vasculature. The model simulates vessel occlusion and pruning due to compression by neighbouring tumour cells; and growth of new vessels in response to hypoxia. Incorporating pressure-mediated vessel occlusion into our ABM required development of a friction force representing cell-extracellular matrix (ECM) adhesion. Through numerical simulations, we show that when friction is neglected, pressure gradients dissipate throughout the tissue and vessel compression is negligible. By contrast, as the strength of the friction force increases, larger pressure gradients form, leading to significant vessel compression.

We perform extensive model simulations, to investigate how changes in the oxygen landscape caused by vascular remodelling affect the tumour’s growth dynamics and composition. We show further how vessel remodelling influences a tumour’s response to radiotherapy. We find tumour regrowth to be especially sensitive to vessel remodelling, with certain vascular landscapes growing rapidly post-radiotherapy, and accelerating tumour regrowth.

## 1 Introduction

Tumours contain a diverse collection of cell populations and cytokines, which, with the extracellular matrix, make up the tumour microenvironment (TME). TME composition varies between tumours, and its evolution is shaped by the constituent tumour cells, infiltrating immune cells, stromal cells, blood vessels and extracellular matrix [1–3]. Interactions between these species produce distinct TME landscapes which can have strong pro-tumour effects [4]. In this paper we develop a multiscale model to investigate the effect that mechanical interactions between proliferating tumour cells and the surrounding vasculature have on oxygen supply to the TME, the tumour’s growth dynamics, and its response to radiotherapy [5].

Tumour growth impacts blood vessels in two important ways. First, mechanical stress generated by rapid tumour growth increases the pressure exerted by tumour cells on blood vessels, causing them to become compressed and occluded, and reducing their blood flow [6, 7]. Further, if the flow rate in a vessel remains sufficiently low for a sufficiently long period then the vessel may be pruned from the vascular network [8]. These factors limit the ability of the vasculature to supply oxygen to the surrounding tissue, leading to reduced oxygen levels which slow tumour growth [9]. This leads to the second effect, where low oxygen levels (hypoxia) stimulate tumour cells to produce diffusible factors such as vascular endothelial growth factor (VEGF) which promote the growth of new blood vessels through angiogenesis [10], one of Hanahan and Weinberg’s hallmarks of cancer [11–13]. The newly formed vessels are typically abnormal, and the associated blood flow is often unstable and highly irregular, which limits their ability to meet the tumour’s oxygen requirements [14, 15].

In this paper, we develop a spatially-resolved model to investigate the effect that interactions between tumour cells and blood vessels have on their co-evolution. Spatially resolved mathematical models of tumour growth fall into two broad categories: continuum models, which describe the time evolution of the tumour density, and agent-based models (ABMs), which distinguish individual cells and whose evolution is governed by pre-defined rules. ABMs can be divided into on-lattice approaches, including cellular automata [16–18] and cellular Potts models [19–21]; and off-lattice approaches such as node-based models [22] and vertex models [23–25]. Hybrid models combine multiple approaches,for example treating certain species or regions as discrete agents (often cells) and others as continuous variables (often diffusible species). For reviews of these different modelling approaches, see [26–31].

We propose a hybrid model, which uses an off-lattice, node-based framework to resolve individual tumour cells and blood vessels and uses a reaction-diffusion equation to determine how the spatial distribution of oxygen evolves over time. This hybrid approach has been used in other models of tumour growth because of its flexibility (see, e.g., [22, 32–40]). Our model builds upon these approaches by accounting for vessel compression caused by mechanical pressure applied to the vessel wall, which causes deformation, and drives vessel occlusion and pruning [6, 7]. Existing ABMs that account for vessel occlusion typically focus on the wall shear stress experienced by blood vessels during blood flow and assume that the wall shear stress must exceed a threshold value if a given vessel is to remain viable and support blood flow: a vessel becomes occluded if the wall shear stress it experiences remains below the threshold value for a sufficiently long period [41–48]. This approach does not account for vessel remodelling due to mechanical pressure generated by tumour growth, an effect shown to be important in experiments [49] and continuum models [50–54].

A key feature of our ABM is the treatment of mechanical interactions between cancer cells and component of the surrounding TME, such as the extracellular matrix (ECM) and extracellular fluid. We account for these interactions with two distinct forces. The first is Stokes’ drag, widely used in ABMs to represent resistance to cell movement due to extracellular fluid. The second, a novel friction force, accounts for resistance to motion due to the continual breaking and formation of cell-ECM bonds of adhesion. While other models account for resistance to motion due to cell-ECM bonds of adhesions [55–57] by explicitly modelling the ECM and its deformation during cell movement, our approach instead models these effects as a combination of friction and drag. Through numerical simulation we demonstrate that friction is necessary for the formation of pressure gradients sufficiently large to drive vessel occlusion: when Stokes’ drag dominates, pressure gradients dissipate across the tissue and vessel compression is negligible. By contrast, as the strength of the friction force increases, larger pressure gradients form, generating significant vessel compression.

We incorporate vessels into the ABM by building on earlier work in which blood vessels were represented as fixed point sources of oxygen, of negligible volume [32, 33, 38, 58]. Here we view vessels as space-occupying agents which interact with tumour cells. While vessel locations are fixed, deformation of vessel walls is caused by forces exerted on them by tumour cells; put simply, the pressure difference across a vessel wall determines whether it is compressed (external pressure exceeds internal pressure) or expands (external pressure less than internal pressure). We assume further that changes in vessel cross-sectional area modulate the supply of oxygen to the tissue and the rate at which vessels are pruned from the system. We incorporate additional rules to account for the growth of new vessels in response to hypoxia.

We demonstrate the impact of including vascular remodelling and the friction force in the ABM by using it to simulate treatment with radiotherapy, a front-line treatment for cancer [59, 60]. We show that the additional modelling assumptions in our ABM can have a significant effect on the tumour’s response to radiotherapy. Due to its prevalence in the clinic, there are many mathematical models of tumour responses to radiotherapy (for example, [61–66]). We follow [58, 62, 67], by simulating radiotherapy with the linear quadratic model. We show that even this simple implementation of radiotherapy, when integrated within our ABM, produces a complex treatment and recovery landscape.

Quantifying the intricate spatial structures generated by ABMs requires the application of spatial analysis methods. Here we use spatial, and shape metrics to describe and quantify a tumour’s vascularisation and morphology. Spatial statistics are commonly used in fields such as astrophysics and ecology and are ideally suited to describe, quantify and compare the location of individual cells in tissue images and ABMs [68–72]. These methods enable us to understand how a tumour and its vasculature evolve over time and as parameters relating to friction and vascular efficacy vary.

The remainder of this paper is structured as follows: in the next section, we summarise our model, focusing on how it extends previous work by Bull & Byrne [32, 33]. We then describe our simulation protocol and parameter sweeps, before introducing the metrics we use to quantify simulation outputs. In the Results section, we demonstrate the importance of friction in enabling pressure to accumulate within a growing tumour and show that such behaviour cannot be generated when cell-ECM interactions are represented by drag forces alone. We investigate further the effect of pressure accumulation on the vasculature, showing how the strength of the friction force and vessel robustness to mechanical stress modulate the degree of vascularisation of a tumour. Vascular remodelling due to tumour growth in turn alters the oxygen landscape, and we show that if the tumour’s demand for oxygen exceeds the supply then this may drive morphological changes that result in multilobular tumours. Finally, we show that mechanically-mediated vascular remodelling affects tumour sensitivity to radiotherapy and the tumour’s subsequent recovery dynamics.

## 2 Methods

In this section we outline our agent-based model (ABM), which builds on an earlier model developed by us [33]. Here we focus on the new elements of the model (full model details are included in S1). We then describe our simulation and parameter sweep protocol, and introduce the methods used to characterise and quantify the qualitative behaviours that the ABM generates.

### 2.1 Agent-Based Model

Our ABM is developed within CHASTE (Cancer, Heart and Soft Tissue Environment), an open source C++ framework for simulating complex, multiphysics, and multiscale mathematical models of biomedical systems, including cancer [73–75]. We adopt a hybrid approach to develop our 2D ABM, using an off-lattice, cell-centre based model to resolve individual tumour cells located within a dynamic, vascular tissue. Following [33], each cell possesses a subcellular cell-cycle model, which determines whether and when a cell proliferates. Proliferation depends on oxygen (*ω*), which is viewed as a diffusible species whose spatial distribution is modelled by a reaction-diffusion equation (S1.1.1), with vessels as point sources and tumour cells as point sinks. As shown in Fig 1A, under normoxic conditions (*ω* ≥ *ω*_*h*_) tumour cells consume oxygen and proliferate. Under low oxygen conditions (*ω*_*n*_ *< ω* ≤ *ω*_*h*_) they stop proliferating while continuing to consume oxygen (S1.1.2). If the local oxygen concentration becomes too low (*ω ≤ ω*_*n*_), then the tumour cells become necrotic and die (via a process described in S1.1.3). Tumour cells also stop proliferating if they are mechanically compressed, i.e., if their area is less than a proportion *η o*f their target area. Cells are physically represented by their centroid and mechanical interactions with nearby cells are modelled via a system of linear springs that connect the cell centroids. Each spring has a target length and resistance which represent each cell’s uncompressed diameter and compressibility respectively. Cells exert forces on their neighbours in order to maintain their target spring lengths (S1.3.1). Fig 1B shows how the balance of forces applied by cells and vessels acting on a cell is resolved and drives cell movement (S1.3.2). For a detailed description, see S1.

**Fig. 1:**
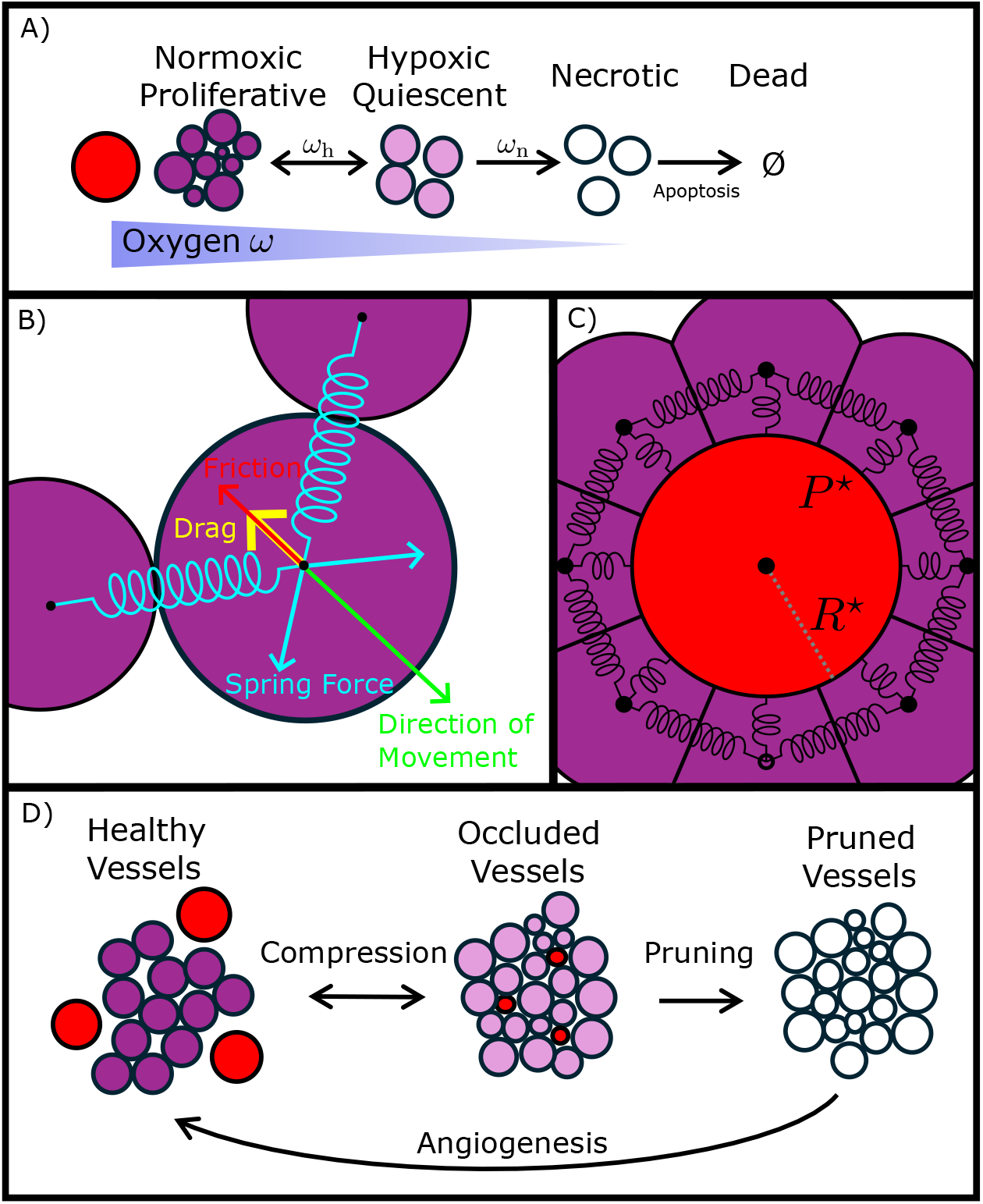
Schematic summarising the key mechanisms in the agent-based model. (A) Tumour cells (purple) consume oxygen which is supplied by blood vessels (red). Tumour cell behaviour is determined by the local oxygen concentration: ‘proliferative’ in well-oxygenated areas (*ω*_*h*_ *< ω*), ‘quiescent’ in intermediate oxygen levels (*ω*_*n*_ *< ω* ≤ *ω*_*h*_) and ‘necrotic’ when oxygen is so low that tumour cells die due to lack of oxygen (0 ≤ *ω* ≤ *ω*_*n*_). (B) Cell movement is determined by balancing the forces acting on a cell. These include spring forces due to physical contact with neighbouring cells which drive movement, and a combination of friction and drag forces which resist movement. (C) Blood vessels experience forces, and in turn apply them to neighbouring tumour cells. The total pressure applied to a vessel is compared to its internal vessel pressure *P* ^⋆^. This pressure difference determines the dynamics of the vessel’s radius *R*^⋆^(*t*), allowing a vessel to be mechanically compressed or to expand. (D) Vessel phenotype is determined by its radius, *R*^⋆^(*t*) which depends on the local cell pressure it experiences: in low pressure the vessel is ‘healthy’. Under compression, it becomes ‘occluded’, and its oxygen supply decreases. If a vessel remains occluded for longer than *τ*_prune_, then it is pruned, and no longer acts as an oxygen source. The lack of oxygen stimulates angiogenesis, whereby a new vessel may form at the same location to restore the oxygen supply.

We now describe the new features of our ABM: vessel remodelling, including occlusion, pruning and angiogenesis; and a friction force which enables pressure to accumulate within the tumour.

#### 2.1.1 Vessel Remodelling

In [33], blood vessels are modelled as fixed point sources of oxygen which do not occupy space. Here, we instead represent them as dynamic agents which occupy space. As before, for simplicity, vessels are assumed to be perpendicular to the plane and, therefore, we represent them as circular agents whose radii evolve in response to environmental cues.

Below, we describe how a vessels supply oxygen to the tissue at varying rates depending on their radius. Then we describe the different phenotypes a blood vessel may adopt and how transitions between phenotypes account for vessel pruning and angiogenesis. Later we explain the force balance which determines the evolution of a vessel’s radius in response to applied forces, see Section 2.1.3.

##### Dynamic Oxygen Supply

A vessel’s radius *R*^⋆^ determines its rate of oxygen supply to the surrounding tissue. The oxygen distribution *ω i*s modelled via a reaction diffusion equation of the form

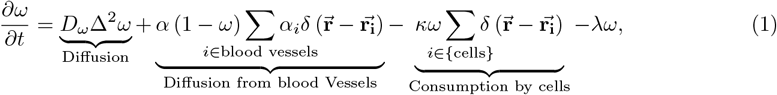

where the positive parameters *D*_*ω*_ and *κ r*epresent the oxygen diffusion coefficient and the rate at which cancer cells consume oxygen respectively. The decay rate *λ* accounts for oxygen consumption by cells not explicitly included in the model. In Equation (1), we assume that the oxygen concentration in the blood vessels is constant, normalised to unity, and that the rate at which the *i*-th blood vessel supplies oxygen to the environment is proportional to the difference between the oxygen concentrations in the blood vessel and the tissue, and denote by *α* the maximum rate of oxygen supply from a blood vessel. We introduce a scaling factor *α*_*i*_ to account for the dependence of the supply rate on the vessel radius *R*_*i*_. For simplicity, we assume that *α*_*i*_ ∈ [0, 1] is a linearly increasing function of *R*_*i*_ of the form

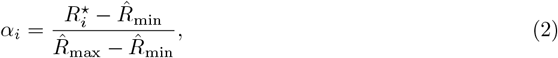

where the parameters 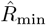 and 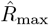 denote the minimum and maximum radii of a blood vessel. We close Equation (1) by imposing no-flux boundary conditions on the domain boundaries:

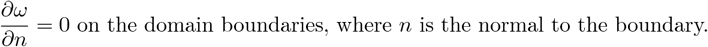

We assume that initially 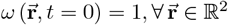.

##### Occlusion, Pruning and Angiogenesis

We distinguish three vessel phenotypes: **Healthy, Occluded** and **Pruned**. The **Healthy** and **Occluded** phenotypes are determined based on the vessel radius:

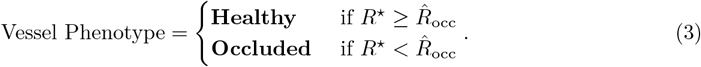

where 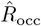 is the threshold vessel radius at which a vessel becomes occluded.

**Healthy** and **Occluded** phenotypes behave similarly, both allow a vessel to supply oxygen to the tissue dependent on the vessel’s radius as described above. Vessels can also quickly transition between **Healthy** and **Occluded** phenotypes as their radius dynamically evolves due to applied forces (see Section 2.1.3). We distinguish the **Occluded** phenotype as vessels which are sufficiently compressed such that they are candidates to be pruned. This is modelled as a vessel’s phenotype switching to **Pruned** if they remain **Occluded** for longer than *τ*_prune_.

**Pruned** vessels are irreparably damaged, considered dead and have been removed from the simulation. **Pruned** vessels may only return to the **Healthy** phenotype following successful angiogenesis. We model angiogenesis by returning **Pruned** vessels to the **Healthy** phenotype under two conditions: First, there must be at least one **Healthy** vessel within distance *D*_*a*ngio_ of the **Pruned** vessel from which the new vessel can emerge. Secondly, the candidate location must be sufficiently hypoxic for an extended period of time. This is defined as *ω < ω*_*a*ngio_ for longer than *τ*_*a*ngio_ and represents the time taken for tumour cells, stimulated by hypoxia, to produce angiogenic factors such as VEGF which stimulate vessel growth. If these two conditions are met then the vessel has probability of *P*_*a*ngio_ to regrow each hour. This is summarised by the pseudocode in S1.2.3 Algorithm 1. The overall system of vessel phenotypes and the transitions between them are summarised in Fig 1D.

#### 2.1.2 Tumour Cell Force Balance

In existing cell-centre models the equations of motion are derived by balancing the applied forces 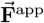 acting on a cell with a Stokes’ drag force 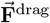, which models the net effect of TME interactions that resist cell movement. Details on the applied contact force 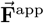 sare given in S1.3.1. In practice, a component of a cell’s interaction with the ECM is the formation of adhesive bonds. These attachments anchor a cell in place and prevent it from moving when small forces are applied; the cell only moves when the applied forces 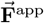 are sufficiently large to break the attachments. Stokes’ drag does not account for this effect, which we model with a friction force 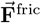.

If 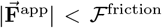 then the applied force is exactly balanced by friction 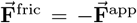, and the cell does not move. In this case, the friction force corresponds to a static friction force and the parameter ℱ^friction^ represents the magnitude of the limiting friction force. If 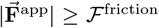 then the friction force becomes dynamic, with fixed magnitude ℱ ^friction^, acting opposite to 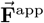. Biologically, the dynamic friction force is the resistance to motion due to continual formation and breaking of cell-ECM attachments. Here we assume that cell-ECM attachments break and form on timescales which are much faster the timescale of cell movement.

Combining these forces, we write the classical equation of motion (4) as follows:

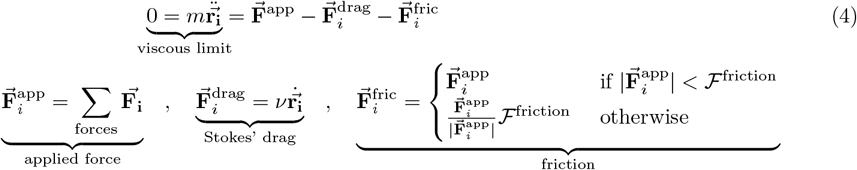

where 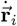 is the velocity of cell *i, ν* is the damping coefficient for Stokes’ drag, and 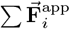 represents the total force acting on cell *i*, due to physical contact with its neighbours.

By neglecting inertial terms (over-damped, viscous limit), Equation (4) can be rearranged to give:

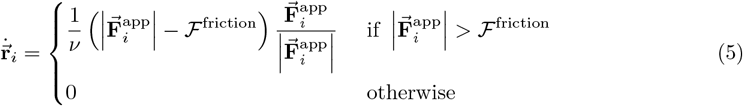

Equation (5) differs from the equation of motion typically used in node-based models: it allows cells to resist applied forces and remain static if the magnitude of the applied force 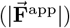 is less than the threshold value ℱ ^friction^.

#### 2.1.3 Vessel Force Balance

Vessel occlusion arises when proliferation of neighbouring tumour cells generates mechanical forces that compresses and occludes vessels. Similarly to tumour cells as described above, we use force balance to derive equations of motion for the vessel due to applied contact forces (details on these contact forces given in S1.3.1). Different to tumour cells, vessels in our model do not move and instead we model the applied forces as acting on and deforming the vessel wall. We model deformations as being driven by a difference in external applied pressure *P*_*i*_, and an internal vessel pressure *P* ^⋆^.

As mentioned above we model vessels by their circular cross-sections with dynamic radii, and we further assume that the mean force exerted by cells surrounding a blood vessel approximates the external pressure *P*_*i*_ applied to a blood vessel. Precisely, letting *N b*e the number of cells neighbouring a given vessel and 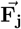 be the force applied by neighbour *j (j =* 1, …, *N)* to the vessel, we can write the pressure contribution *ρ*_*j*_ from neighbour *j a*s:

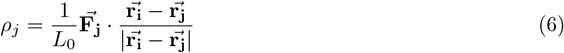

where *L*_*0*_ is the length of the cell-vessel interface over which contact pressure is applied,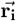 is the vessel’s position and 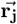 is the position of neighbour *j*. We approximate each cell-vessel interface to have length *L*_*0*_ = 1 since cell boundaries are poorly defined in the overlapping-spheres framework we are using. We also assume that each neighbour’s pressure contribution *ρ*_*j*_ is evenly distributed along the vessel boundary and therefore assume that the pressure *P*_*i*_ applied to a vessel is the average of the pressure contributions from each neighbour:

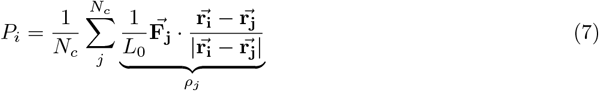

Equation (7) defines how we model the mechanical pressure experienced by a vessel due to intercellular forces. We acknowledge that alternative methods could be used to approximate mechanical pressure, but consider this simple functional form to be appropriate here given uncertainties in how the modelled spring forces translate into physical contact forces.

The difference between the external pressure *P*_*i*_ (7) and internal pressure *P* ^⋆^ generates a force of magnitude *L*_*0*_ (*P*_*i*_ − *P* ^⋆^) acting to occlude (*P*_*i*_ *> P* ^⋆^) or dilate (*P*_*i*_ *< P* ^⋆^) the vessel. By balancing this against a damping force with damping coefficinet *ν*_*r*_ and assuming that the maximum vessel radius is 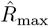, we derive an equation of motion for the vessel’s radius *R*_*i*_ = *R*_*i*_ (*t)*:

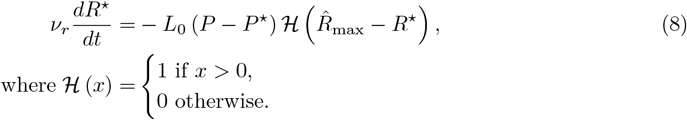

##### 2.1.4 Radiotherapy

Our model generates heterogeneous oxygen landscapes due to interactions between the tumour cells and the surrounding vasculature. We demonstrate the impact of this heterogeneity by simulating tumour responses to radiotherapy using a simple model of cell death following exposure to radiotherapy. We focus on the cell killing due to radiotherapy as it is known to be sensitive to oxygen levels [65, 66].

We adapt a simple Linear-Quadratic (LQ) model [62] to describe the probability that a tumour cell dies following exposure to a single dose of radiotherapy, *P (***death**|dose). This probability depends on the radiotherapy dose, the tumour’s oxygen status and the local oxygen concentration:

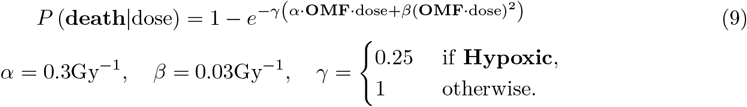

*γ a*ccounts for reduced radio-sensitivity of quiescent, hypoxic cells due to their lack of DNA replication [61], whilst the Oxygen Modification Factor (**OMF**) accounts for reduced effectiveness of radiotherapy due to the lack of oxygen needed to generate DNA damaging free radicals [63]:

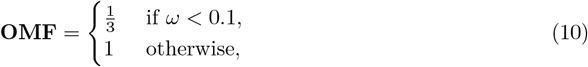

where *ω* is the local oxygen concentration.

In radiotherapy simulations, treatment is applied at *t =* 21 days and simulations continued for a further 21 days. Any cells killed by radiotherapy are labelled as ‘apoptotic’ and die in the same way as necrotic cells (for further details, S1.1.3).

### 2.2 Simulating Tumours

Each simulation starts with 4 tumour cells seeded at the centre of a 1mm by 1mm square domain. 400 ‘healthy’ blood vessels and 600 ‘pruned’ vessels are randomly placed in the domain without overlap, representing a well vascularised tissue. The randomness in the initial vessel distribution is distinct from other stochastic parts of the model and can be varied using a configuration seed parameter which is separate from the random seed used in the rest of the model. This allows stochastic simulations to be conducted in the same vascular environment if desired. Synthetic tumours grow for 42 days of simulation time or until the number of tumour cells exceeded 10,000. Snapshots of the tumour are saved at 10 hour timesteps.

We analyse the model’s qualitative behaviour by performing four parameter sweeps: two focused parameter sweeps in the absence of radiotherapy, one in the presence of radiotherapy, and an extensive Latin hypercube sampled parameter sweep in the absence of radiotherapy which we use to test the robustness of findings from the focused parameter sweeps. The parameter sweeps are summarised in Table 1.

**Table 1:** Summary of parameter sweeps. Table summarising parameter sweeps performed to analyse model. *ℛ*: Whether radiotherapy was applied at day 21 *N*_*p*_: Number of parameter sets in parameter sweep *N*_*c*_: Number of initial vessel configurations simulated *N*_*r*_ : Number of stochastic repetitions Total: Total number of simulations in parameter sweep

| Parameter Sweep | $\mathcal{R}$ | Parameters Varied | $N_p$ | $N_c$ | $N_r$ | Total | Notes |
| --- | --- | --- | --- | --- | --- | --- | --- |
| $P^\star - \mathcal{F}^{\text{friction}}$ | No | $\mathcal{F}^{\text{friction}}, P^\star$ | 110 | 4 | 4 | **1,760** | S2.1.1 |
| $\omega_h - P^\star$ | No | $\omega_h, P^\star$ | 70 | 4 | 4 | **1,120** | S2.1.2 |
| Radiotherapy | Yes | $P^\star, \omega_{\text{angio}}$ | 100 | 4 | 4 | **1,600** | S2.2 |
| Latin Hypercube | No | $\kappa, \omega_h, P^\star, \mathcal{F}^{\text{friction}}, \eta, \hat{R}_{\text{occ}}, \nu, \mu$ | 5000 | 4 | 1 | **20,000** | S2.1.3 |
| **Total** |  |  |  |  |  | **24,480** |  |

### 2.3 Spatial analysis

#### 2.3.1 Tumour-Vessel Pair Correlation Function

We use the cross pair correlation function (cross-PCF) to quantify the spatial distribution of blood vessels as they are remodelled by the tumour [71, 72, 76]. The tumour-vessel cross-PCF (*g*_*BT*_ (*r)*) measures the number of observed tumour-vessel pairs separated by distance *r*, relative to the number of such pairs expected under a null hypothesis of complete spatial randomness (CSR). We define annuli with inner radii *r a*nd outer radii *r + dr*, centred around each blood vessel, and consider how many tumour cells fall within these annuli (see Fig 2A). Letting 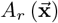 be the area of the intersection of an annulus of radius *r c*entred at 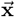 and the simulation domain, and defining an indicator function

**Fig. 2:**
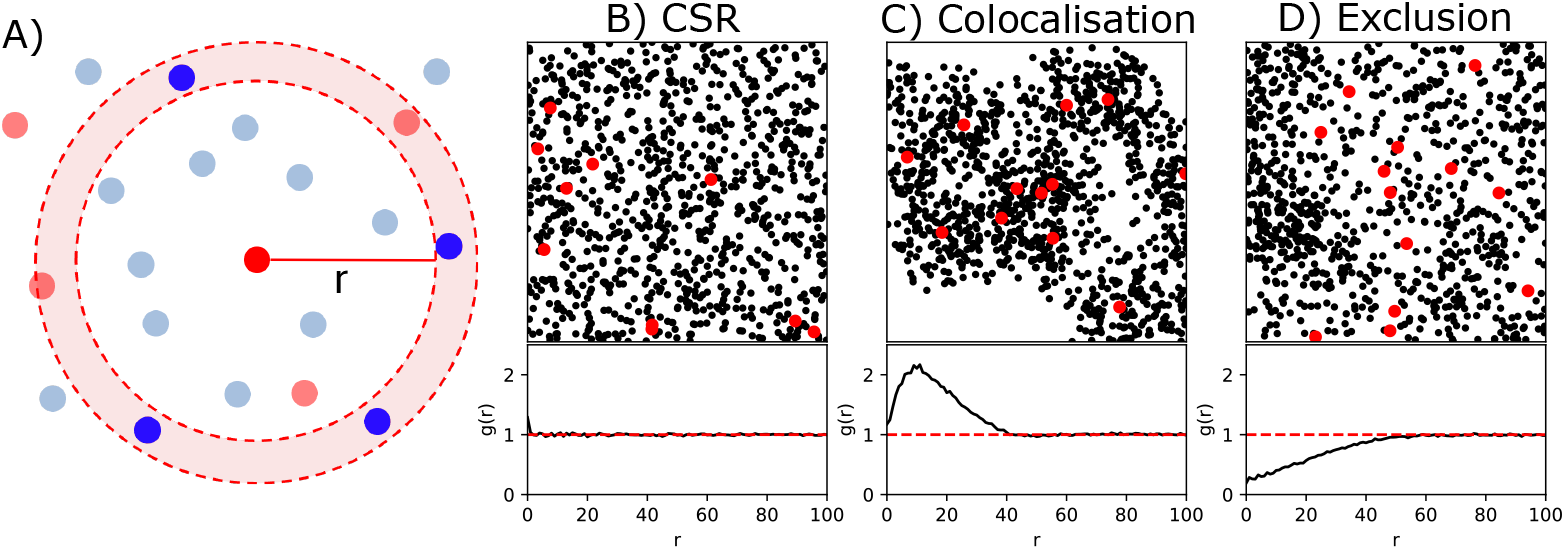
Schematic of cross pair correlation function. (A) Schematic demonstrating how the cross-PCF is calculated by drawing annuli around red points and counting the number of blue points which fall within the annuli. (B, C, D) Illustrations of the cross-PCF applied to synthetic datasets. Letting red points represent vessels and black points represent tumour cells, we demonstrate three basic spatial signatures. (B) Complete spatial randomness (CSR). The cross-PCF is flat (*g*_*BT*_ (*r*) ≃ 1). (C) Colocalisation of vessels and tumour cells. The cross-PCF has a peak whose height indicates the strength of colocalisation and whose *r* value indicates the range of interaction distances at which the two agent types are positively correlated. (D) Negative correlation between vessels and tumour cell location. Here, *g*_*BT*_ *<* 1 for *r <* 50, indicating exclusion of tumour and vessels points at this length scale.

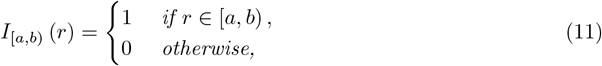

the tumour-vessel PCF is defined as

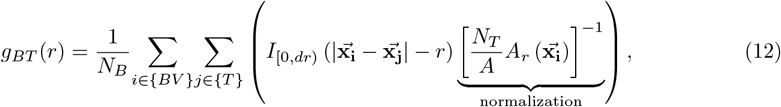

where {*BV* } are blood vessels, {*T* } are tumour cells, *N*_*B*_ and *N*_*T*_ are the numbers of blood vessels and tumour cells respectively. Here, *g*_*BT*_ (*r)* > 1 represents colocalisation of tumour cells and blood vessels at radius *r*, whereas *g*_*BT*_ (*r)* < 1 indicates exclusion. Practically, the cross-PCF defined in Equation 12 is calculated at a discrete series of radii *r*_*k*_ such that *r*_*0*_ = 0, *r*_*k*+1_ = *r*_*k*_ + *dr*. We fix *dr =* 1 cell diameter and consider radii up to a maximum of *r =* 100 cell diameters.

To aid interpretation of the cross-PCF, Figs 2B,C,D show three synthetic datasets, involving red (representing vessels) and black (representing tumour cells) points, and their cross-PCF signatures. Under complete spatial randomness (CSR), with no spatial correlation between red and black points (Fig 2B), the cross-PCF is flat and close to 1 for all *r*. When black points cluster around red points (Fig 2C), there is a peak in the cross-PCF; its height (*g(*10) ≃ 2.2) quantifies the increase in pairs observed separated by this distance compared to the number expected under CSR, and its location (*r*≃ 10) indicates the length scale at which clustering is strongest. Such cross-PCF signatures indicate increased tumour-vessel colocalisation at short distances which we interpret as the presence of vessels within the main tumour mass. In Fig 2D, black points are sparse around red points, and the resultant PCF has *g(r)* < 1 for *r* < 50. Such cross-PCF signatures indicate exclusion of blood vessels from the tumour.

The cross-PCF for small *r* will be important in our analysis because it indicates the local colocalisation, or exclusion, of blood vessels and tumour cells. Therefore, we define the mean cross-PCF value for 0 *< r* ≤ 5 cell diameters, denoted 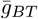, as:

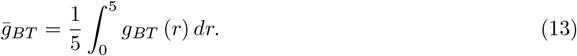

#### 2.3.2 Tumour Roundness

To quantify a simulated tumour’s morphology, we apply a simple metric which describes the tumour’s roundness. First, we determine the tumour’s bounding polygon by calculating its *α-*shape [77] with critical parameter 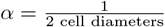 . The roundness of this polygon is defined as:

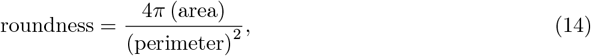

which is 1 for a circle and tends to 0 as the elongation of an oval goes towards infinity. While roundness could be quantified in other ways [78], including eccentricity and roughness [79], we use this metric because it is simple and interpretable, and captures changes in tumour morphology which provide insight into tumour behaviour.

### 2.4 Response to Radiotherapy

The radiotherapy model described in 2.1.4 determines which cells are killed. To quantify its effects, we use two metrics. First, we measure χ, the percentage of tumour cells that die during a simulated dose of radiation. Secondly, we record *T*_*R*_, the time it takes for the tumour to regrow to the same number of cells as immediately before the radiation dose was applied. Denoting by 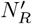 and *N*_*R*_ the number of cells killed and the number of live cells immediately before radiotherapy, we define 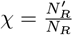. The recovery time, *T*_*R*_, is the time taken for the number of tumour cells to return to *N*_*R*_. As stated in 2.1.4, simulations are continued for 21 simulated days after treatment, by which time all tumours had regrown to at least *N*_*R*_ cells.

To understand the behaviour of *T*_*R*_ it is useful to study how the tissue’s vasculature evolves before and after radiotherapy. Particularly relevant is Ω, the oxygen capacity of the vasculature, which is defined as follows:

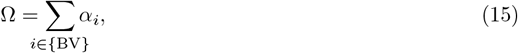

where {BV} are blood vessels, and the scaling factor *α*_*i*_ determines vessel *i’s abili*ty to supply oxygen as defined in equation (2).

## 3 Results

In this section, we show that including a friction force in our off-lattice ABM enables pressure to accumulate within a tumour. We then investigate how pressure accumulation impacts blood vessel occlusion, and show how dynamic interactions between blood vessels and tumour cells affect a tumour’s growth dynamics and its morphology. Finally, we show how vascular remodelling impacts tumour responses to radiotherapy.

### 3.1 Simulating a friction force enables accumulation of pressure

We first consider a simple model in which a compressive force is applied at the left hand boundary of a 1D chain of 100 cells. Specific details of this toy model are given in S1.4. We compare the pressure that accumulates when there is no friction force (Fig 3A), and when a friction force is included (Fig 3B).

**Fig. 3:**
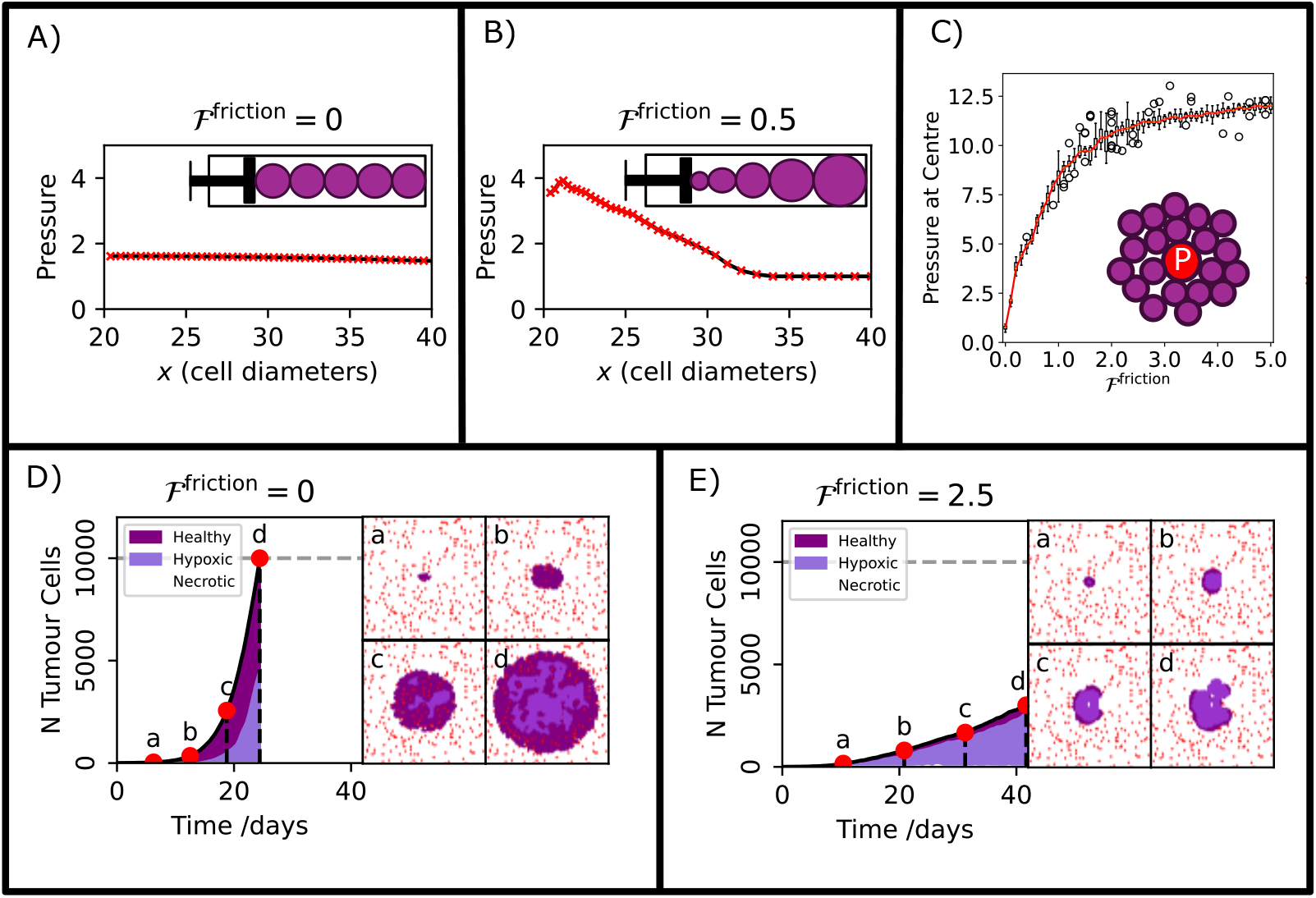
Friction drives pressure accumulation inside tumours. (A) 1D simulation in which a chain of cells is compressed by slowly moving the left boundary from *x* = 0 to *x* = 20, in the absence of friction (ℱ^friction^ = 0.0). At equilibrium, all cells experience the same pressure. (B) The simulation in A) is repeated with *F* ^*friction*^ = 0.5 (all other parameters held fixed). At equilibrium, cells closer to the applied force experience higher pressure than those farther away. In contrast to a), the force has no impact on cells at distance from the applied force. (C) Tumour cells surrounding a single blood vessel grow until the tumour reaches an equilibrium size. The equilibrium pressure on the blood vessel, averaged over 16 repeats, is plotted as a function of the friction strength, demonstrating that without friction, pressure does not accumulate in the centre of the cluster. (D,E) Tumour growth dynamics exhibited by our model in the absence (D) and presence (E) of a friction force, with all other model parameters fixed at the default values specified in S1.5. Snapshots a-d show the simulation at four highlighted timepoints; red: blood vessels, purple: healthy tumour, lilac: hypoxic tumour, white: necrotic tumour. (D) Simulation without friction (ℱ ^friction^ = 0), (E) Simulation including friction (ℱ^friction^ = 2.5). Without friction, blood vessels withstand the pressure due to tumour growth, resulting in high tissue oxygenation and a large, rapidly growing tumour. When friction is included, pressure accumulates causing blood vessel occlusion, the formation of a hypoxic core inside the tumour, and slower tumour growth.

The pressure distributions plotted in Figs 3A and B show that incorporating friction permits the accumulation of pressure within the chain of cells. In the absence of friction (Fig 3A), the compressive force is transmitted evenly to all cells in the chain, creating constant pressure. When friction is included (Fig 3B), pressure accumulates within cells which are close to the compressed boundary on the left, and decreases with distance from the boundary, eventually reaching a constant value, so that distant cells do not move in response to the applied force.

In a second example, the growth of a population of tumour cells is supported by a single blood vessel which is unaffected by pressure and, as such, cannot be occluded. In this case, the tumour grows around the vessel until its reaches a stable, equilibrium size. Fig 3C shows how the value of the friction strength ℱ^friction^ affects the compressive pressure acting on the blood vessel at the centre of the tumour when it reaches its equilibrium size. The pressure that accumulates at the centre of the tumour is an increasing, saturating function of ℱ^friction^, demonstrating that the inclusion of friction causes the pressure associated with tumour growth to accumulate.

These simple examples illustrate how frictional forces may impact the distribution of pressure in a simulated tissue, and in Figs 3D,E we show that this can impact tumour growth dynamics. Figs 3D,E show typical simulations in which a tumour is seeded in a vascular environment, in the absence (D) and presence (E) of a friction force. In Fig 3D, we fix ℱ^friction^ = 0. The pressure disperses uniformly across the cells and does not accumulate inside the tumour. As a result, the blood vessels remain functional and a large tumour, containing approximately 10, 000 cells, forms after approximately 23 days. By contrast, Fig 3E shows that when ℱ^*friction*^ > 0 (all other model parameters held fixed), pressure accumulates inside the tumour, vessels become occluded, leading to the formation of a hypoxic core and slowing tumour growth.

### 3.2 Pressure accumulation causes vessel occlusion and formation of avascular tumours

We use the tumour-vessel cross-PCF, g_*BT*_ (r), to quantify the spatial distribution of blood vessels within a tumour and, in so doing, evaluate the degree of vessel remodelling.

Fig 4A shows how *g*_*BT*_ (r) can be used to distinguish vascular and avascular tumours. We consider two simulations, generated using the same parameter values, except for *P* ^⋆^ (high *P* ^⋆^ left, low *P* ^⋆^ right), and show the final simulation timepoints alongside the corresponding tumour-vessel cross-PCFs. For high *P* ^⋆^, the blood vessels do not experience sufficient pressure to be occluded, and, instead, dense, well vascularised cell clusters surround them. The colocalisation of tumour cells and blood vessels is captured in the cross-PCF, with *g*_*BT*_ (r) > 1 for r < 25. Conversely, for low *P* ^⋆^, the accumulated pressure exceeds the vessel pressure, causing vessel occlusion and the formation of an avascular tumour. The absence (or exclusion) of vessels from the tumour is captured in the cross-PCF, with *g*_*BT*_ < 1 for r < 25.

**Fig. 4:**
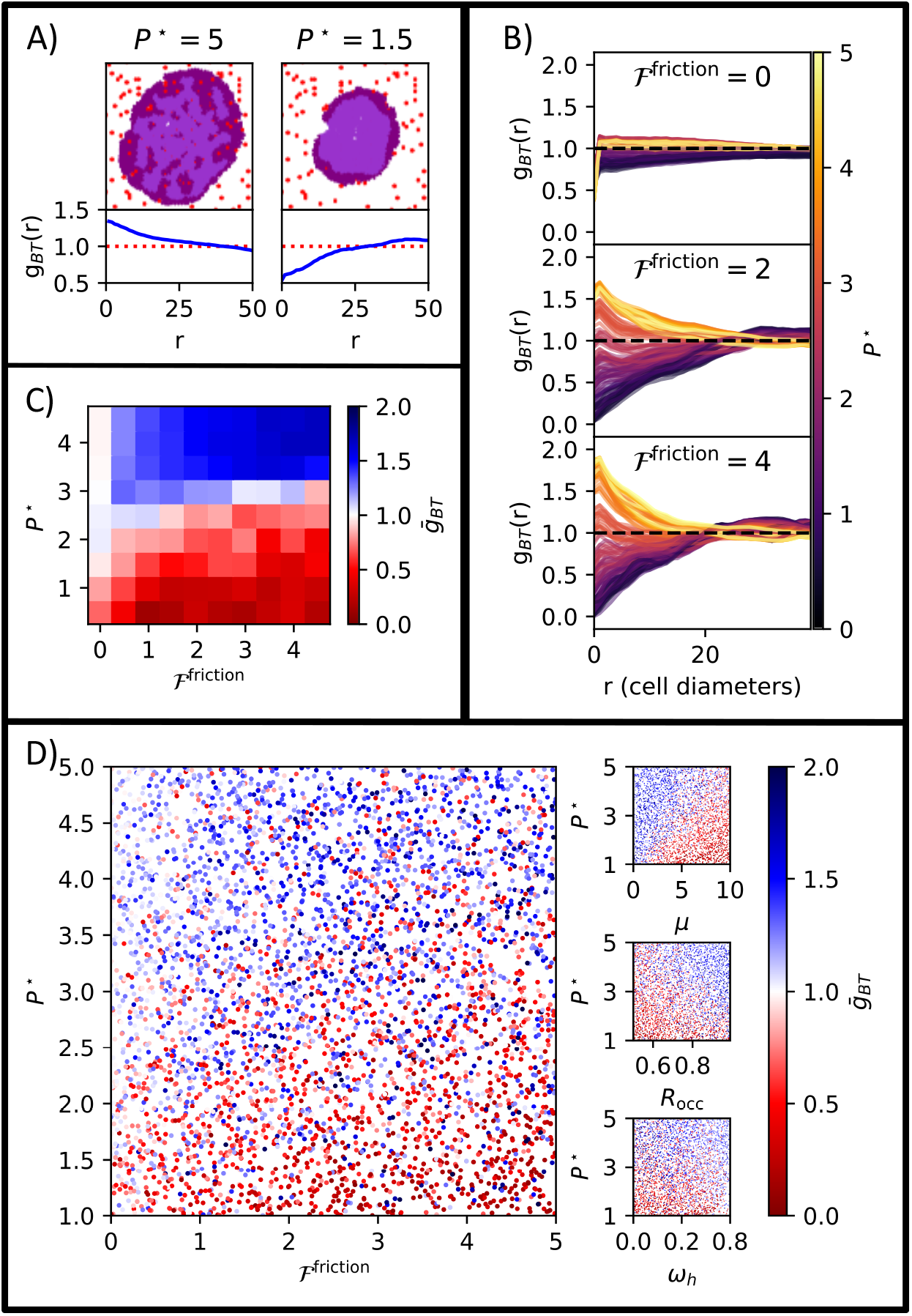
Pressure accumulation drives vessel occlusion and emergence of avascular tumours. (A) Tumour distributions and tumour-vessel cross-PCFs, *g*_*BT*_ (*r*), from two representative simulations with different values of *P* ^⋆^ (*t* = 28 days). Left: High *P* ^⋆^ = 5 leads to a well-vascularised tumour, with colocalisation of tumour cells and vessels (*g*_*BT*_ *>* 1 for *r <* 25). Right: Low *P* ^⋆^ = 1.5 leads to the formation of an avascular tumour, which lacks vessels due to occlusion (*g*_*BT*_ *<* 1 for *r <* 25). Red: blood vessels, purple: normoxic tumour, lilac: hypoxic tumour. For this example ℱ^friction^ = 1 and other parameters are default values given in S1.5. (B) Tumour-Vessel cross-PCF signatures for fixed ℱ^friction^ coloured by *P* ^⋆^ (S2.1.1), demonstrating that, when *F*^friction^ *>* 0, the value of *P* ^⋆^ can substantially impact tumour vascularisation (characterised by *g*_*BT*_ signatures that are qualitatively similar to those for the vascular and avascular tumours shown in (A). For ℱ^friction^ = 0, all tumours are well-vascularised, for all values of *P* ^⋆^. For 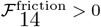, small values of *P* ^⋆^ lead to formation of avascular tumours and larger *P* ^⋆^ values lead to well-vascularised tumours. (C) 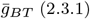, a metric measuring the low (red) or high (blue) vascularisation of the tumour across the *P* ^⋆^ ℱ^friction^ 2-parameter sweep in S2.1.1. There is a threshold value of *P* ^⋆^, which depends on ℱ^friction^, below which tumours become avascular. (D) 2-parameter projections of 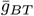 across a multidimensional latin hypercube parameter sweep (S2.1.3). Main panel shows 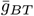 as a function of *P* ^⋆^ and ℱ^friction^, and shows a similar trend to panel C but with some noise due to changes in other model parameters. Sub panels show how, in the same simulations, 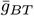 varies with three additional parameters; cell stiffness *µ*, vessel occlusion 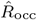 and tumour hypoxia sensitivity *ω*_*h*_. The values of these parameters impact 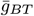, suggesting they contribute to whether a tumour remainsvascular.

Fig 4B shows the g_*BT*_ (r) signature associated with the final simulation timepoints from a 2-parameter sweep over ℱ^friction^ and P ^⋆^ (S2.1.1). When ℱ^friction^ > 0 (middle and bottom plots of Fig 4B) the cross-PCFs are heterogenous, and qualitative comparison with the *g*_*BT*_ signatures in 4A indicates that the tumours range from well vascularised to avascular. When ℱ^friction^ > 0, the value of P ^⋆^ determines how the *g*_*BT*_ signatures transition from those associated with well vascularised tumours in which cells cluster around vessels to those associated with avascular tumours in which vessels have been occluded by tumour growth. In contrast, when ℱ^friction^ = 0 (uppermost panel of Fig 4B), the cross-PCF signatures remain relatively flat for all values of P ^⋆^. In these cases, the tumour grows alongside the vessels with pressure quickly dissipating as cells are not anchored to the substrate. This both restricts the formation of dense clusters around vessels and prevents vessel occlusion, resulting in flatter g_*BT*_ signatures. These results highlight the importance of friction in our model: ℱ^friction^ allows cells to anchor to the substrate and prevents the dispersion of local mitotic forces, which drives the accumulation of pressure. The magnitude of the accumulated pressure relative to P ^⋆^ determines whether vessels become occluded, and explains why the dependence of vascularisation (as characterised by g_*BT*_) on P ^⋆^ disappears when ℱ^friction^ = 0; pressure cannot accumulate when ℱ^friction^ = 0.

Using the metric 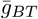 to quantify vascularisation as ℱ^friction^ and *P* ^⋆^ vary reinforces this interpretation. The results in Fig 4C show that low values of *P* ^⋆^ result in 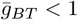 (avascular tumours), whilst high values result in 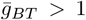 (vascular tumours). The transition between 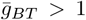 and 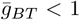 is pronounced when ℱ^friction^ > 0 and becomes less pronounced as ℱ^friction^ →0. We note further from Fig 4C that ℱ^friction^ defines the threshold value of *P* ^⋆^ at which a tumour switches from being vascular to avascular.

In order to determine the robustness of the trends seen in this 2-parameter sweep, we performed a larger, Latin-hypercube parameter sweep, varying 8 parameters (S2.1.3). Fig 4D shows a projection of the Latin hypercube parameter sweep in which *P* ^⋆^ and ^friction^ range over the same values as in panel C. We observe the same qualitative trends, with reduced *P* ^⋆^ resulting in vessel occlusion, and the value of ℱ^friction^ determining the *P* ^⋆^ threshold at which this occurs. However the relationship is noisier, indicating that other model parameters influence vessel exclusion.

Analysis of alternative projections in our Latin hypercube parameter sweep reveal 3 additional parameters which impact vessel exclusion (see sub panels in Fig 4D). These parameters relate to a cell’s ability to generate mechanical forces which can occlude vessels (*µ*), a blood vessel’s resistance to remodelling 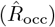 and a tumour cell’s oxygen requirements (ω_*h*_).

### 3.3 The oxygen landscape affects tumour morphology

In Section 3.2 we saw how the value of the vessel pressure *P* ^⋆^ determines whether a tumour’s vessels are occluded. Since blood vessels supply oxygen, varying *P* ^⋆^ also alters the oxygen landscape. In this section, we perform a 2-parameter sweep (S2.1.2) to determine how varying *P* ^⋆^ and *ω*_*h*_ affects a tumour’s growth dynamics (recall that *ω*_*h*_ is the threshold oxygen concentration below which cells halt proliferation and become quiescent).

First, we show how changing ω_*h*_ can affect tumour size, composition and morphology at *t* = 42 days. In Fig 5A, the tumour on the left has a lower *ω*_*h*_ value. As a result, its cells can withstand lower oxygen concentrations and remain viable at greater distances from blood vessels, leading to a compact tumour mass. The tumour on the right has a higher *ω*_*h*_ value which limits its growth to small lobes that surround blood vessels.

**Fig. 5:**
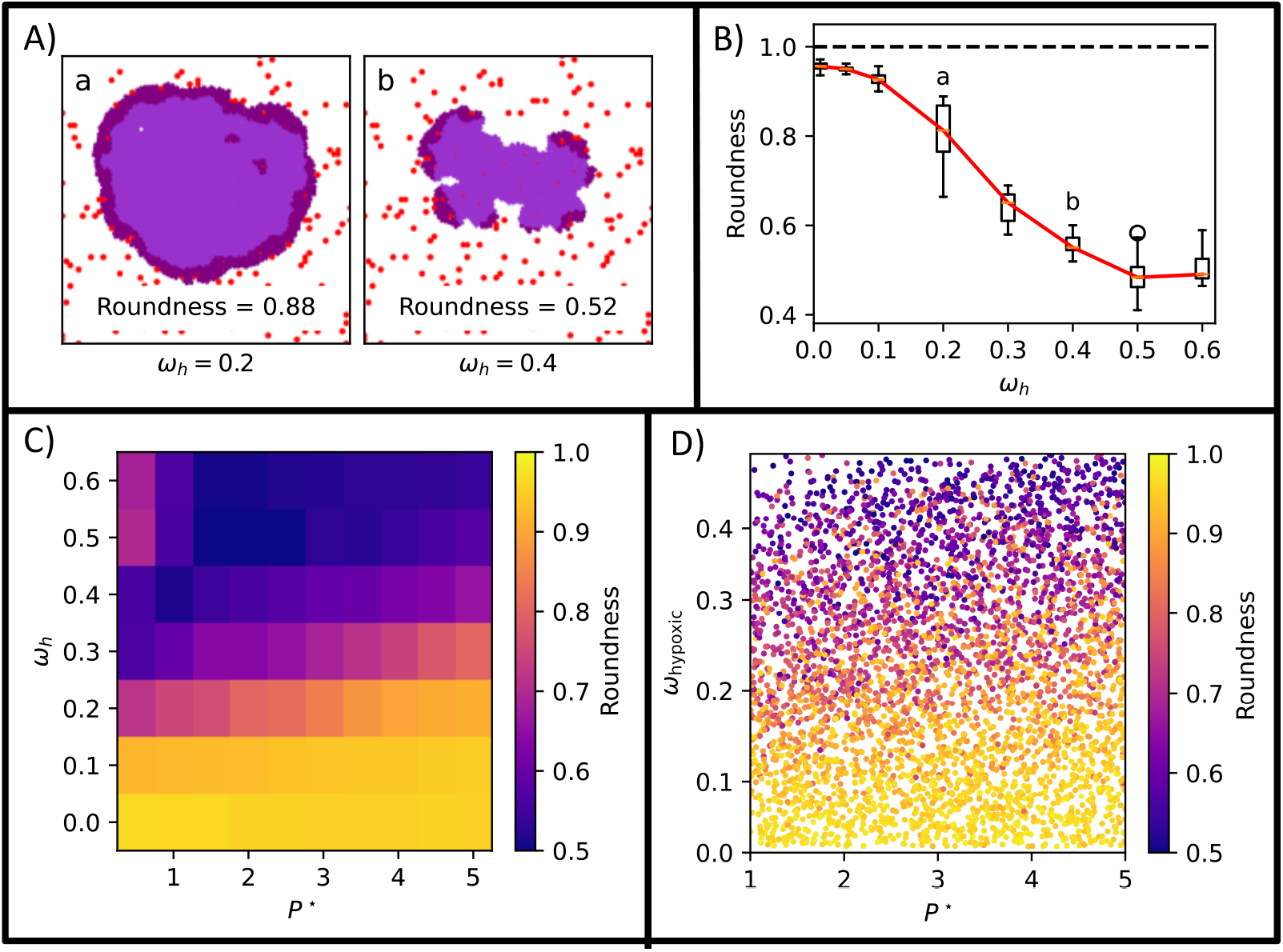
The oxygen landscape drives the formation of tumour lobes. Tumour roundness is calculated at the end timepoint for each simulation. (A) Representative tumours at *t* = 42 days from the *P* ^⋆^ − *ω*_*h*_ parameter sweep (S2.1.2) showing how tumour morphology becomes more lobular as *ω*_*h*_ increases from *ω*_*h*_ = 0.2 (Left) to *ω*_*h*_ = 0.4 (Right), and how this can be captured by the roundness score. Red: blood vessel, purple: normoxic tumour, lilac: hypoxic tumour. (B) Effect of varying *ω*_*h*_ on tumour roundness when *P* ^⋆^ = 2, showing a sigmoidal relationship between *ω*_*h*_ (oxygen requirement) and the extent to which the tumour forms lobes. Markers a and b indicate the parameter values for the tumours shown in panel A. (C) Heatmap showing how roundness changes as *P* ^⋆^ and *ω*_*h*_ vary (for other parameters see S2.1.2). Roundness scores are averaged across 4 realisations for each parameter set and show that smaller values of *P* ^⋆^ shift the sigmoidal relationship in panel B towards smaller values of *ω*_*h*_. (D) 2-dimensional projection from our multidimensional Latin hypercube sweep(S2.1.3) onto the same axes as C, demonstrating that the behaviour in our 2-parameter sweep is robust to variations in other parameters in the multidimensional parameter sweep.

We quantify tumour morphology using the roundness score (see Section 2.3.2), observing that roundness decreases as ω_*h*_ increases and the tumour becomes more lobular. Fig 5B shows how tumour roundness decreases in a sigmoidal manner as *ω*_*h*_ increases, indicating that as oxygen demand increases, the tumour forms lobes and becomes less round. Fig 5C shows how tumour roundness changes as *P* ^⋆^ and ω_*h*_ vary. As in Fig 5B, the roundness score decreases as *ω*_h_ increases. Varying P ^⋆^ shifts the sigmoid-like dependence on ω_*h*_ in Fig 5B: for smaller *P* ^⋆^, the roundness score decreases at smaller *ω*_*h*_. The observed change in roundness can be understood as the tumour being restricted to lobes that surround blood vessels when the oxygen supply (low *P* ^⋆^) is insufficient for the oxygen requirement (*ω*_*h*_). We performed similar analyses to investigate how tumour mass and hypoxic fraction vary across parameter space. As expected, increased vascularisation resulted in larger and better oxygenated tumours (see S3.1 for details).

As before, we performed a larger Latin hypercube parameter sweep (S2.1.3) to determine whether the trends for the roundness score persist when multiple model parameters vary. In Fig 5D we project results from the larger parameter sweep onto (P ^⋆^, ω_*h*_) parameter space. The trends from Fig 5C persist, suggesting that the changes in tumour morphology described above are more sensitive to P ^⋆^ and ω_*h*_ than other parameters in the Latin hypercube (S2.1.3).

### 3.4 Vascular remodelling impacts tumour response to radiotherapy

In this section we use our model to investigate how vessel remodelling impacts a tumour’s response to radiotherapy. We employ the simplified oxygen-dependent radiotherapy model and radiotherapy protocol described in Section 2.1.4: a single dose of radiotherapy is applied at t = 21 days after which the tumour grows for another 21 days. All parameters are fixed at default values (S1.5), except for the vessel parameters P ^⋆^ and ω_angio_ which vary as described in S2.2.

Fig 6A shows representative simulated tumours that were treated with radiotherapy on day 21. Parameters are given in S2.2. The vasculature for Tumour (a) has a large value of P ^⋆^ and is, therefore, large, well oxygenated and sensitive to radiotherapy. Tumour (c) has a low value of P ^⋆^. As a result, its vessels are easily occluded and the oxygen supply is low, leading to the formation of a radio-resistant, hypoxic core. The value of P ^⋆^ for Tumour (b) and its response to radiotherapy are intermediate between those for Tumours (a) and (c). Whilst its immediate response to radiotherapy appears qualitatively similar to Tumour (a), its post radiotherapy growth is substantially different.

**Fig. 6:**
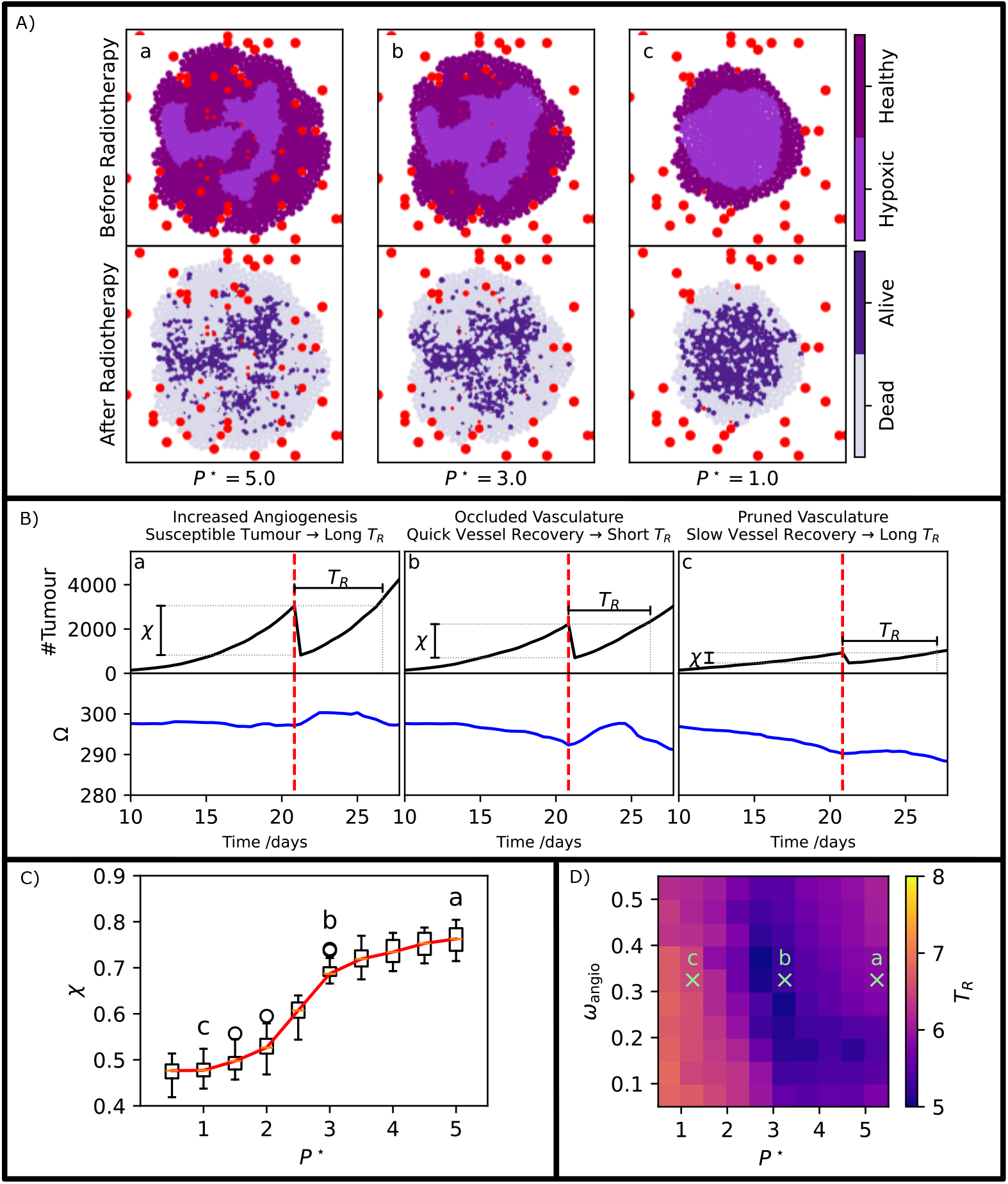
Effect of dynamic vasculature on radiotherapy outcomes. (A) Representative simulations showing tumours immediately before and after exposure to radiotherapy for different values of *P* ^⋆^. Large values of *P* ^⋆^ (*P* ^⋆^ = 5.0 in ‘a’) give rise to well oxygenated tumour which are extremely sensitive to radiotherapy so that a large proportion of the tumour cells are killed by radiotherapy. As *P* ^⋆^ decreases (*P* ^⋆^ = 3.0 in ‘b’ and *P* ^⋆^ = 1.0 in ‘c’), the tumour forms a radio-resistant, hypoxic core which enables a larger proportion of tumour cells to survive treatment. For these tumours, *ω*_angio_ = 0.3 and other parameters are fixed at the default values given in S1.5. (B) Tumour and blood vessel growth trajectories for the simulations shown in a. The percentage of tumour cells killed (𝒳) decreases as *P* ^⋆^ decreases, reinforcing the results from panel A where low *P* ^⋆^ creates radiotherapy resistance. The parameter Ω, defined in 2.4, measures the vasculature’s oxygen capacity. (C) Variation in *χ* as *P* ^⋆^ varies. At low *P* ^⋆^ vessel occlusion causes hypoxia and radiotherapy resistance. As *P* ^⋆^ increases, both vascularisation and tissue oxygenation increase, and we identify a threshold value of *P* ^⋆^ at which tumours switch from low to high oxygenation. (D) Change in the post-radiotherapy recovery time *T*_*R*_ as *P* ^⋆^ and *ω*_angio_ vary. A valley of fast recovery times (‘b’) extends from a region with low *P* ^⋆^ and high *ω*_angio_ to a region with high *P* ^⋆^ and low *ω*_angio_.

The growth trajectories of the tumour and vasculature for Tumours (a), (b) and (c) are plotted in Fig 6B. We use the percentage of tumour killed (𝒳) and post-radiotherapy recovery time (T_*R*_), described in Section 2.4, to quantify the tumour’s response to radiotherapy. We observe that 𝒳 decreases as P ^⋆^ decreases from Tumour (a) to (c), reinforcing our interpretation of Fig 6A above. Fig 6C shows how 𝒳 varies with P ^⋆^ across a slice of our parameter sweep (S2.2) with ω_angio_ = 0.3. These results further support the relationship between 𝒳 and P ^⋆^ described above. Additionally, Fig 6C shows that there is a threshold P ^⋆^ (P ^⋆^ ≃ 2.5 for the chosen parameter regime) below which 𝒳 falls significantly due to the formation of a large hypoxic core. 𝒳 is strongly dependent on P ^⋆^, with ω_angio_ having a much smaller effect (for further details, see S3.2).

Fig 6B suggests that the dependence of T_*R*_ on P ^⋆^ differs from that for 𝒳, with T_*R*_ being smallest for Tumour (b). Whilst we might expect Tumour (a), with high P ^⋆^, to have a long T_*R*_ because it is highly radiosensitive, it is initially less clear why Tumour (c) also has a long recovery time. We can understand this by looking at the dynamics of Ω, the oxygen capacity of the vasculature (see Section 2.4). For Tumour (b), the vasculature recovers rapidly following radiotherapy, and T_*R*_ is short.

For Tumour (c), the vasculature recovers more slowly, resulting in increased T_*R*_. This difference in vessel recovery depends on the condition of the vasculature pre-radiotherapy. For Tumour (b), P ^⋆^ is sufficiently small that vessels are readily occluded, causing hypoxia and conferring radiotherapy resistance. However, the value of P ^⋆^ is not small enough for vascular pruning. By contrast, for Tumour (c), P ^⋆^ is small enough for vascular pruning. The resulting hypoxia leads to radiotherapy resistance, and necessitates angiogenesis to replace pruned vessels post-radiotherapy, greatly slowing tumour recovery.

We investigate this behaviour further in Fig 6D, where we show how T_*R*_ changes as P ^⋆^ and ω_angio_ vary. We observe a a valley of fast recovery times which includes Tumour (b) and extends from low P ^⋆^ and high ω_*angio*_ to high P ^⋆^ and low ω_*angio*_. In order to understand this landscape, we note that two competing effects contribute to the length of the recovery time T_*R*_. First, as P ^⋆^ and ω_angio_ increase, tumour cells are better oxygenated and, hence, more susceptible to radiotherapy, which causes 𝒳 to increase. As a result, T_*R*_ also increases because more tumour cells must proliferate to replace those removed by radiotherapy: this explains why Tumour (a) is characterised by high 𝒳 and T_*R*_. Secondly, for sufficiently small values of P ^⋆^ the vasculature is heavily pruned following radiotherapy. Here, even though the tumour is radiotherapy resistant due to poor oxygenation, the slow rate of vessel regrowth limits the overall rate of tumour regrowth, increasing the recovery time, T_*R*_. This explains why Tumour (c) has low 𝒳 and high T_*R*_. The valley of short recovery times forms where P ^⋆^ is low enough for poor oxygenation to confer radiotherapy resistance, but high enough that the vasculature is not pruned. Instead, the vasculature is occluded and, therefore, the death of tumour cells post-radiotherapy reduces the mechanical pressure on the vessels, allowing them to quickly recover and increase the supply of oxygen, enabling the tumour cells to regrow quickly. This explains why Tumour (b) recovers faster than Tumours (a) and (c).

## 4 Discussion

We have developed an off-lattice agent-based model that describes the growth of a population of tumour cells embedded within a vascular tissue. Our ABM accounts for mechanical interactions between the tumour cells and blood vessels and vascular remodelling. We used our model to study the effect that the dynamic oxygen landscape associated with vascular remodelling has on a tumour’s morphology and response to radiotherapy.

Our model accounts for the growth of new vessels stimulated by tumour hypoxia, as well as their occlusion and removal due to tissue stress caused by rapid tumour growth. The mechanical components of off-lattice ABMs typically operate in the non-inertial, viscous limit and introduce a resistive force (representing cell-substrate interactions) to balance external forces acting on individual cells. Most existing off-lattice ABMs model this resistive force via Stokes’ drag. In such models, pressure dissipates over the tissue domain due to the dependence of Stokes’ drag on the velocity. As a result, forces generated by proliferating cells are quickly transmitted to the tumour boundary, preventing tumour cells from generating sufficient mechanical pressure to occlude blood vessels. In order to enable vessel occlusion, we introduced into the mechanical model a friction force which accounts for the adhesive bonds that form between the cells and the extracellular matrix and prevents a cell from moving if the applied forces it experiences are below a threshold value. Our simulations show that including this friction force, together with Stokes’ drag, prevents rapid pressure dissipation and enables more physically realistic modelling of vessel occlusion.

In addition to vessel occlusion and pruning caused by rapid tumour growth, our vascular model accounts for the growth of new vessels in regions that experience prolonged hypoxia. We demonstrated that, in combination with friction, *P* ^⋆^ (the pressure within a vessel) determines how readily vessels are occluded and pruned from the tumour. In more detail, increased *P* ^⋆^ results in well vascularised tumours and decreased *P* ^⋆^ results in avascular tumours.

We showed further how changes in the vasculature may impact tumour morphology. We quantified how increased sensitivity of tumour cells to hypoxia (high *ω*_*h*_) can cause a tumour to form lobes that encircle blood vessels and become less round if there is insufficient oxygen supply (low *P* ^⋆^). In the supporting information, we demonstrate that changes to the vasculature may impact other tumour features, including its size and hypoxic fraction (S3.1). These results illustrate the importance of considering vascular occlusion when modelling tumour growth. Our model also highlights the importance of accounting for friction in force-based models of tumour growth, in order to enable pressure to accumulate within the tissue and to occlude blood vessels.

Finally, we showed that the addition of friction and vessel remodelling can have a significant effect on tumour responses to radiotherapy. The vessel pressure parameter *P* ^⋆^ affects the percentage of the tumour killed by radiation, with high *P* ^⋆^ generating normoxic tumours that are susceptible to radiotherapy, and low *P* ^⋆^ resulting in hypoxic tumours which are radioresistant, supporting observations in earlier studies [58, 64]. Tumours within which vessels have very high *P* ^⋆^ have increased recovery time. As *P* ^⋆^ decreases the recovery time decreases, matching the reduced sensitivity to radiotherapy. However, there is a threshold *P* ^⋆^ below which the vasculature becomes both severely occluded and, importantly, pruned. This leads to significantly increased recovery times, as tumour recovery is then limited by the slower process of vessel angiogenesis. These results highlight the substantial impact that vessel mechanisms can have on a tumour’s response to radiotherapy.

There are many ways in which we could extend the work presented in this paper. We could relax our simplified view of the vasculature as a series of point sources of oxygen. Following [41, 45, 80–82], we could instead represent it as a dynamic network of connected vessels and model blood flow and oxygen transport through the network. Given the importance of cell-ECM interactions in generating mechanical stress within solid tumours [83, 84], and motivated by existing models of cell-ECM interactions [85–88] (reviewed in [55, 89]), it would also be interesting in future work to incorporate the ECM into our model, to explicitly model the formation and breaking of cell-ECM adhesion bonds and study their impact on cell movement and ECM deformation. It would also be interesting to investigate the effect of vessel occlusion on other forms of therapy such as immunotherapy [90] and chemotherapy [91–93], which have been studied in similar vascular models focusing on angiogenesis [37, 44, 45]. The design principles used to develop our ABM mean that it is ideally suited to study the complex mechanisms involved in these treatments and their combinations. Future work could also compare our results against experimental data. Cutting-edge multiplex imaging methods [94] can locate and phenotype individual cell in a sample, leading to the prospect of future data becoming available which matches the spatial resolution of our model. The methods we have used to analyse our simulated data could be applied to such imaging data [68–70], leading to the potential for model parametrisation to be conducted from imaging data.

In summary, we have used our model to demonstrate that vessel remodelling, especially pressure-mediated vessel occlusion, can have a significant effect on both tumour growth and sensitivity to radiotherapy. In developing our model, we have also identified possible limitations of existing off-lattice models and shown how they can be resolved by incorporating a friction force, highlighting the need for more work to investigate how cell-substrate interactions can be incorporated into cell-based models of biological tissues, including vascular tumours.

## Data Availability Statement

All code and data associated with this paper can be found at https://github.com/NicholasFan235/VascularRemodelling_Fan2025.

## Acknowledgments

NF was funded by Engineering and Physical Sciences Research Council (EPSRC) grant number EP/S024093/1; National Institute for Health and Care Research (NIHR) Oxford Biomedical Research Centre (BRC) grant HPR00322; and Cancer Research UK (CRUK).

JAB was supported by Cancer Research UK (CRUK) grant number CTRQQR-2021 100002, through the Cancer Research UK Oxford Centre.

## References

[1] Mina J. Bissell and Derek Radisky. “Putting tumours in context”. In: Nature Reviews Cancer 1.1 (Oct. 2001). Publisher: Nature Publishing Group, pp. 46–54. ISSN: 1474-1768. doi: 10.1038/35094059. URL: https://www.nature.com/articles/35094059 (visited on 12/18/2024).

[2] Shelly Maman and Isaac P. Witz. “A history of exploring cancer in context”. In:Nature Reviews Cancer 18.6 (June 2018). Publisher: Nature Publishing Group, pp. 359–376. ISSN: 1474-1768. doi:10.1038/s41568-018-0006-7. URL: https://www.nature.com/articles/s41568-018-0006-7 (visited on 12/18/2024).

[3] Nicole M. Anderson and M. Celeste Simon. “The tumor microenvironment”. In:Current biology: CB 30.16 (Aug. 17, 2020), R921–R925. ISSN: 1879-0445. doi:10.1016/j.cub.2020.06.081.

[4] Alicia Cristina Peña-Romero and Esteban Orenes-Piñero. “Dual Effect of Immune Cells within Tumour Microenvironment: Pro- and Anti-Tumour Effects and Their Triggers”. In:Cancers 14.7 (Jan. 2022). Number: 7 Publisher: Multidisciplinary Digital Publishing Institute, p. 1681. ISSN: 2072-6694. doi:10.3390/cancers14071681. URL: https://www.mdpi.com/2072-6694/14/7/1681 (visited on 12/18/2024).

[5] Melissa R. Junttila and Frederic J. de Sauvage. “Influence of tumour micro-environment heterogeneity on therapeutic response”. In:Nature 501.7467 (Sept. 2013). Publisher: Nature Publishing Group, pp. 346–354. ISSN: 1476-4687. doi:10.1038/nature12626. URL: https://www.nature.com/articles/nature12626 (visited on 12/18/2024).

[6] Timothy P. Padera et al. “Cancer cells compress intratumour vessels”. In:Nature 427.6976 (Feb. 2004). Publisher: Nature Publishing Group, pp. 695–695. ISSN: 1476-4687. doi:10.1038/427695a. URL: https://www.nature.com/articles/427695a (visited on 12/13/2024).

[7] Genevièeve Griffon-Etienne et al. “Taxane-induced Apoptosis Decompresses Blood Vessels and Lowers Interstitial Fluid Pressure in Solid Tumors: Clinical Implications1”. In:Cancer Research 59.15 (Aug. 1, 1999), pp. 3776–3782. ISSN: 0008-5472.

[8] Claudia Korn and Hellmut G. Augustin. “Mechanisms of Vessel Pruning and Regression”. In:Developmental Cell 34.1 (July 6, 2015), pp. 5–17. ISSN: 1534-5807. doi:10.1016/j.devcel.2015.06.004. URL: https://www.sciencedirect.com/science/article/pii/S1534580715003937 (visited on 12/13/2024).

[9] Rakesh K. Jain, John D. Martin, and Triantafyllos Stylianopoulos. “The Role of Mechanical Forces in Tumor Growth and Therapy”. In:Annual Review of Biomedical Engineering 16 (Volume 16, 2014 July 11, 2014). Publisher: Annual Reviews, pp. 321–346. ISSN: 1523-9829, 1545-4274. doi:10.1146/annurev-bioeng-071813-105259. URL: https://www.annualreviews.org/content/journals/10.1146/annurev-bioeng-071813-105259 (visited on 12/18/2024).

[10] L. Claesson-Welsh and M. Welsh. “VEGFA and tumour angiogenesis”. In:Journal of Internal Medicine 273.2 (2013). eprint: https://onlinelibrary.wiley.com/doi/pdf/10.1111/joim.12019, pp. 114–127. ISSN: 1365-2796. doi:10.1111/joim.12019. URL: https://onlinelibrary.iley.com/doi/abs/10.1111/joim.12019 (visited on 12/18/2024).

[11] Douglas Hanahan and Robert A. Weinberg. “The Hallmarks of Cancer”. In:Cell 100.1 (Jan. 7, 2000). Publisher: Elsevier, pp. 57–70. ISSN: 0092-8674, 1097-4172. doi:10.1016/S0092-8674(00)81683-9. URL: https://www.cell.com/cell/abstract/S0092-8674(00)81683-9 (visited on 12/18/2024).

[12] Douglas Hanahan and Robert A. Weinberg. “Hallmarks of Cancer: The Next Generation”. In: Cell 144.5 (Mar. 4, 2011), pp. 646–674. ISSN: 0092-8674, 1097-4172. doi:10.1016/j.cell.2011.02.013. URL: https://www.cell.com/cell/abstract/S0092-8674(11)00127-9 (visited on 12/12/2024).

[13] Douglas Hanahan. “Hallmarks of Cancer: New Dimensions”. In: Cancer Discovery 12.1 (Jan. 12, 2022), pp. 31–46. ISSN: 2159-8274. doi: 10.1158/2159-8290.CD-21-1059. URL: https://doi.org/10.1158/2159-8290.CD-21-1059 (visited on 12/12/2024).

[14] Roberta Lugano, Mohanraj Ramachandran, and Anna Dimberg. “Tumor angiogenesis: causes, consequences, challenges and opportunities”. In: Cellular and Molecular Life Sciences 77.9 (May 1, 2020), pp. 1745–1770. ISSN: 1420-9071. doi: 10.1007/s00018-019-03351-7. URL: https://doi.org/10.1007/s00018-019-03351-7 (visited on 12/18/2024).

[15] Rakesh K. Jain. “Determinants of Tumor Blood Flow: A Review1”. In: Cancer Research 48.10 (May 1, 1988), pp. 2641–2658. ISSN: 0008-5472.

[16] AALPEN A. Patel et al. “A Cellular Automaton Model of Early Tumor Growth and Invasion: The Effects of Native Tissue Vascularity and Increased Anaerobic Tumor Metabolism”. In: Journal of Theoretical Biology 213.3 (Dec. 7, 2001), pp. 315–331. ISSN: 0022-5193. doi: 10.1006/jtbi.2001.2385. URL: https://www.sciencedirect.com/science/article/pii/S0022519301923859 (visited on 12/17/2024).

[17] Christian A. Yates, Andrew Parker, and Ruth E. Baker. “Incorporating pushing in exclusion-process models of cell migration”. In: Physical Review E 91.5 (May 22, 2015). Publisher: American Physical Society, p. 052711. doi: 10.1103/PhysRevE.91.052711. URL: https://link.aps.org/doi/10.1103/PhysRevE.91.052711 (visited on 12/17/2024).

[18] Sabine Dormann and Andreas Deutsch. “Modeling of Self-Organized Avascular Tumor Growth with a Hybrid Cellular Automaton”. In: In Silico Biology 2.3 (Jan. 1, 2002). Publisher: IOS Press, pp. 393–406. ISSN: 1386-6338. URL: https://content.iospress.com/articles/in-silico-biology/isb00058 (visited on 12/17/2024).

[19] Marc Durand and Etienne Guesnet. “An efficient Cellular Potts Model algorithm that forbids cell fragmentation”. In: Computer Physics Communications 208 (Nov. 1, 2016), pp. 54–63. ISSN: 0010-4655. doi: 10.1016/j.cpc.2016.07.030. URL: https://www.sciencedirect.com/science/article/pii/S0010465516302284 (visited on 12/17/2024).

[20] François Graner and James A. Glazier. “Simulation of biological cell sorting using a two-dimensional extended Potts model”. In: Physical Review Letters 69.13 (Sept. 28, 1992). Publisher: American Physical Society, pp. 2013–2016. doi: 10.1103/PhysRevLett.69.2013. URL: https://link.aps.org/doi/10.1103/PhysRevLett.69.2013 (visited on 12/17/2024).

[21] STEPHEN Turner and JONATHAN A. Sherratt. “Intercellular Adhesion and Cancer Invasion: A Discrete Simulation Using the Extended Potts Model”. In: Journal of Theoretical Biology 216.1 (May 7, 2002), pp. 85–100. ISSN: 0022-5193. doi: 10.1006/jtbi.2001.2522. URL: https://www.sciencedirect.com/science/article/pii/S0022519301925226 (visited on 12/17/2024).

[22] Ignacio Ramis-Conde et al. “Multi-scale modelling of cancer cell intravasation: the role of cadherins in metastasis”. In: Physical Biology 6.1 (Mar. 2009), p. 016008. ISSN: 1478-3975. doi: 10.1088/1478-3975/6/1/016008. URL: https://dx.doi.org/10.1088/1478-3975/6/1/016008 (visited on 12/05/2024).

[23] Alexander G. Fletcher et al. “Vertex Models of Epithelial Morphogenesis”. In: Biophysical Journal 106.11 (June 3, 2014). Publisher: Elsevier, pp. 2291–2304. ISSN: 0006-3495, 1542-0086. doi: 10.1016/j.bpj.2013.11.4498. URL: https://www.cell.com/biophysj/abstract/S0006-3495(13)05794-9 (visited on 12/17/2024).

[24] Alexander G. Fletcher et al. “Implementing vertex dynamics models of cell populations in biology within a consistent computational framework”. In: Progress in Biophysics and Molecular Biology 113.2 (Nov. 1, 2013), pp. 299–326. ISSN: 0079-6107. doi: 10.1016/j.pbiomolbio.2013.09.003. URL: https://www.sciencedirect.com/science/article/pii/S0079610713000989 (visited on 12/17/2024).

[25] Reza Farhadifar et al. “The Influence of Cell Mechanics, Cell-Cell Interactions, and Proliferation on Epithelial Packing”. In: Current Biology 17.24 (Dec. 18, 2007). Publisher: Elsevier, pp. 2095–2104. ISSN: 0960-9822. doi: 10.1016/j.cub.2007.11.049. URL: https://www.cell.com/current-biology/abstract/S0960-9822(07)02334-2 (visited on 12/17/2024).

[26] Philipp M. Altrock, Lin L. Liu, and Franziska Michor. “The mathematics of cancer: integrating quantitative models”. In: Nature Reviews Cancer 15.12 (Dec. 2015). Publisher: Nature Publishing Group, pp. 730–745. ISSN: 1474-1768. doi: 10.1038/nrc4029. URL: https://www.nature.com/articles/nrc4029 (visited on 12/18/2024).

[27] James M. Osborne et al. “Comparing individual-based approaches to modelling the selforganization of multicellular tissues”. In: PLOS Computational Biology 13.2 (Feb. 13, 2017). Publisher: Public Library of Science, e1005387. ISSN: 1553-7358. doi: 10.1371/journal.pcbi.1005387. URL: https://journals.plos.org/ploscompbiol/article?id=10.1371/journal.pcbi.1005387 (visited on 12/13/2024).

[28] John Metzcar et al. “A Review of Cell-Based Computational Modeling in Cancer Biology”. In: JCO Clinical Cancer Informatics 3 (Feb. 4, 2019), p. CCI.18.00069. ISSN: 2473-4276. doi: 10.1200/CCI.18.00069. URL: https://www.ncbi.nlm.nih.gov/pmc/articles/PMC6584763/ (visited on 12/18/2024).

[29] T. Alarćon, H. M. Byrne, and P. K. Maini. “Towards whole-organ modelling of tumour growth”. In: Progress in Biophysics and Molecular Biology. Modelling Cellular and Tissue Function 85.2 (June 1, 2004), pp. 451–472. ISSN: 0079-6107. doi: 10.1016/j.pbiomolbio.2004.02.004. URL: https://www.sciencedirect.com/science/article/pii/S007961070400032X (visited on 12/17/2024).

[30] Katarzyna A. Rejniak and Alexander R. A. Anderson. “Hybrid models of tumor growth”. In: WIREs Systems Biology and Medicine 3.1 (2011). eprint: https://onlinelibrary.wiley.com/doi/pdf/10.1002/wsbm pp. 115–125. ISSN: 1939-005X. doi: 10.1002/wsbm.102. URL: https://onlinelibrary.wiley.com/doi/abs/10.1002/wsbm.102 (visited on 12/17/2024).

[31] Tiina Roose, S. Jonathan Chapman, and Philip K. Maini. “Mathematical Models of Avascular Tumor Growth”. In: SIAM Review 49.2 (Jan. 2007), pp. 179–208. ISSN: 0036-1445, 1095-7200. doi: 10.1137/S0036144504446291. URL: http://epubs.siam.org/doi/10.1137/S0036144504446291 (visited on 12/17/2024).

[32] Joshua A. Bull et al. “Mathematical modelling reveals cellular dynamics within tumour spheroids”. In: PLOS Computational Biology 16.8 (Aug. 18, 2020). Publisher: Public Library of Science, e1007961. ISSN: 1553-7358. doi: 10.1371/journal.pcbi.1007961. URL: https://journals.plos.org/ploscompbiol/article?id=10.1371/journal.pcbi.1007961 (visited on 12/17/2024).

[33] Joshua A. Bull and Helen M. Byrne. “Quantification of spatial and phenotypic heterogeneity in an agent-based model of tumour-macrophage interactions”. In: PLOS Computational Biology 19.3 (Mar. 27, 2023), e1010994. ISSN: 1553-7358. doi: 10.1371/journal.pcbi.1010994. URL: https://journals.plos.org/ploscompbiol/article?id=10.1371/journal.pcbi.1010994 (visited on 12/12/2024).

[34] H. L. Rocha et al. “A hybrid three-scale model of tumor growth”. In:Mathematical Models and Methods in Applied Sciences 28.1 (Jan. 2018). Publisher: World Scientific Publishing Co., pp. 61–93. ISSN: 0218-2025. doi:10.1142/S0218202518500021. URL: https://www.worldscientific.com/doi/abs/10.1142/S0218202518500021 (visited on 12/17/2024).

[35] Heber L. Rocha et al. “A persistent invasive phenotype in post-hypoxic tumor cells is revealed by fate mapping and computational modeling”. In:iScience 24.9 (Sept. 24, 2021). Publisher: Elsevier. ISSN: 2589-0042. doi:10.1016/j.isci.2021.102935. URL: https://www.cell.com/iscience/abstract/S2589-0042(21)00903-2 (visited on 12/17/2024).

[36] Inês G. Gonçalves and Jose Manuel Garcia-Aznar. “Extracellular matrix density regulates the formation of tumour spheroids through cell migration”. In:PLOS Computational Biology 17.2 (Feb. 26, 2021). Publisher: Public Library of Science, e1008764. ISSN: 1553-7358. doi: 10.1371/journal.pcbi.1008764. URL: https://journals.plos.org/ploscompbiol/article?id=10.1371/journal.pcbi.1008764 (visited on 12/13/2024).

[37] Tobias Duswald et al. “Bridging scales: A hybrid model to simulate vascular tumor growth and treatment response”. In:Computer Methods in Applied Mechanics and Engineering418 (Jan. 2024), p. 116566. ISSN: 00457825. doi:10.1016/j.cma.2023.116566. URL: https://linkinghub.elsevier.com/retrieve/pii/S0045782523006904 (visited on 12/17/2024).

[38] Jessica L. Kingsley et al. “Bridging cell-scale simulations and radiologic images to explain short-time intratumoral oxygen fluctuations”. In:PLOS Computational Biology 17.7 (July 26, 2021). Publisher: Public Library of Science, e1009206. ISSN: 1553-7358. doi:10.1371/journal.pcbi.1009206. URL: https://journals.plos.org/ploscompbiol/article?id=10.1371/journal.pcbi.1009206 (visited on 12/02/2024).

[39] Cicely K. Macnamara et al. “Computational modelling and simulation of cancer growth and migration within a 3D heterogeneous tissue: The effects of fibre and vascular structure”. In: Journal of Computational Science40 (Feb. 1, 2020), p. 101067. ISSN: 1877-7503. doi:10.1016/j.jocs.2019.101067. URL: https://www.sciencedirect.com/science/article/pii/S1877750319305691 (visited on 12/17/2024).

[40] Caleb M. Phillips et al. “A hybrid model of tumor growth and angiogenesis: In silico experiments”. In:PLOS ONE 15.4 (Apr. 10, 2020). Publisher: Public Library of Science, e0231137. ISSN: 1932-6203. doi:10.1371/journal.pone.0231137. URL: https://journals.plos.org/plosone/article?id=10.1371/journal.pone.0231137 (visited on 12/02/2024).

[41] T. Alarćon, H. M. Byrne, and P. K. Maini. “A Multiple Scale Model for Tumor Growth”. In:Multiscale Modeling & Simulation 3.2 (Jan. 2005). Publisher: Society for Industrial and Applied Mathematics, pp. 440–475. ISSN: 1540-3459. doi:10.1137/040603760. URL: https://epubs.siam.org/doi/abs/10.1137/040603760 (visited on 12/17/2024).

[42] Markus R. Owen et al. “Angiogenesis and vascular remodelling in normal and cancerous tissues”. In:Journal of Mathematical Biology 58.4 (Apr. 1, 2009), pp. 689–721. ISSN: 1432-1416. doi:10.1007/s00285-008-0213-z. URL: https://doi.org/10.1007/s00285-008-0213-z (visited on 12/17/2024).

[43] Holger Perfahl et al. “Multiscale Modelling of Vascular Tumour Growth in 3D: The Roles of Domain Size and Boundary Conditions”. In:PLOS ONE 6.4 (Apr. 13, 2011). Publisher: Public Library of Science, e14790. ISSN: 1932-6203. doi:10.1371/journal.pone.0014790. URL: https://journals.plos.org/plosone/article?id=10.1371/journal.pone.0014790 (visited on 12/18/2024).

[44] Steven R. McDougall, Alexander R. A. Anderson, and Mark A. J. Chaplain. “Mathematical modelling of dynamic adaptive tumour-induced angiogenesis: Clinical implications and therapeutic targeting strategies”. In:Journal of Theoretical Biology 241.3 (Aug. 7, 2006), pp. 564–589. ISSN: 0022-5193. doi:10.1016/j.jtbi.2005.12.022. URL: https://www.sciencedirect.com/science/article/pii/S0022519305005564 (visited on 12/17/2024).

[45] A. Stéphanou et al. “Mathematical modelling of the influence of blood rheological properties upon adaptative tumour-induced angiogenesis”. In:Mathematical and Computer Modelling. Advances in Business Modeling and Decision Technologies [pp. 1-95] 44.1 (July 1, 2006), pp. 96– 123. ISSN: 0895-7177. doi:10.1016/j.mcm.2004.07.021. URL: https://www.sciencedirect.com/science/article/pii/S0895717705004565 (visited on 12/17/2024).

[46] K. Bartha and H. Rieger. “Vascular network remodeling via vessel cooption, regression and growth in tumors”. In:Journal of Theoretical Biology 241.4 (Aug. 2006), pp. 903–918. ISSN: 00225193. doi:10.1016/j.jtbi.2006.01.022. URL: https://linkinghub.elsevier.com/retrieve/pii/S0022519306000373 (visited on 12/15/2024).

[47] Y. Lee et al. “A cellular automaton model for the proliferation of migrating contact-inhibited cells”. In:Biophysical Journal 69.4 (Oct. 1, 1995). Publisher: Elsevier, pp. 1284–1298. ISSN: 0006-3495, 1542-0086. doi:10.1016/S0006-3495(95)79996-9. URL: https://www.cell.com/biophysj/abstract/S0006-3495(95)79996-9 (visited on 12/17/2024).

[48] M. Welter, K. Bartha, and H. Rieger. “Emergent vascular network inhomogeneities and re-sulting bloodflow patterns in a growing tumor”. In:Journal of Theoretical Biology 250.2 (Jan. 2008), pp. 257–280. ISSN: 00225193. doi:10.1016/j.jtbi.2007.09.031. URL: https://linkinghub.elsevier.com/retrieve/pii/S0022519307004584 (visited on 12/15/2024).

[49] Morgan Delarue et al. “Compressive Stress Inhibits Proliferation in Tumor Spheroids through a Volume Limitation”. In:Biophysical Journal 107.8 (Oct. 21, 2014), pp. 1821–1828. ISSN: 0006-3495. doi:10.1016/j.bpj.2014.08.031. URL: https://www.ncbi.nlm.nih.gov/pmc/articles/PMC4213738/ (visited on 02/13/2025).

[50] R. P. Araujo and D. L. S. McElwain. “New insights into vascular collapse and growth dynamics in solid tumors”. In:Journal of Theoretical Biology 228.3 (June 7, 2004), pp. 335–346. ISSN: 0022-5193. doi:10.1016/j.jtbi.2004.01.009. URL: https://www.sciencedirect.com/science/article/pii/S0022519304000384 (visited on 12/17/2024).

[51] R. P. Araujo and D. L. S. McElwain. “The role of mechanical host–tumour interactions in the collapse of tumour blood vessels and tumour growth dynamics”. In:Journal of Theoretical Biology 238.4 (Feb. 21, 2006), pp. 817–827. ISSN: 0022-5193. doi:10.1016/j.jtbi.2005.06.033. URL: https://www.sciencedirect.com/science/article/pii/S0022519305002912 (visited on 12/17/2024).

[52] A.F. Jones et al. “A mathematical model of the stress induced during avascular tumour growth”. In:Journal of Mathematical Biology 40.6 (June 1, 2000), pp. 473–499. ISSN:1432-1416. doi:10.1007/s002850000033. URL: https://doi.org/10.1007/s002850000033 (visited on 12/17/2024).

[53] Min Wu et al. “The effect of interstitial pressure on tumor growth: Coupling with the blood and lymphatic vascular systems”. In:Journal of Theoretical Biology320 (Mar. 7, 2013), pp. 131–151. ISSN: 0022-5193. doi:10.1016/j.jtbi.2012.11.031. URL: https://www.sciencedirect.com/science/article/pii/S0022519312006200 (visited on 12/17/2024).

[54] M. J. Holmes and B. D. Sleeman. “A Mathematical Model of Tumour Angiogenesis Incorporating Cellular Traction and Viscoelastic Effects”. In:Journal of Theoretical Biology 202.2 (Jan. 21, 2000), pp. 95–112. ISSN: 0022-5193. doi:10.1006/jtbi.1999.1038. URL: https://www.sciencedirect.com/science/article/pii/S002251939991038X (visited on 12/17/2024).

[55] Vincent Nöel et al. “PhysiMeSS - a new physiCell addon for extracellular matrix modelling”. In:Gigabyte2024 (Oct. 16, 2024). Publisher: GigaScience Press, gigabyte136. ISSN: 2709-4715. doi:10.46471/gigabyte.136. URL: https://gigabytejournal.com/articles/136 (visited on 12/18/2024).

[56] Gianlu C. D’antonio, Paul Macklin, and Luigi Preziosi. “AN AGENT-BASED MODEL FOR ELASTO-PLASTIC MECHANICAL INTERACTIONS BETWEEN CELLS, BASEMENT MEMBRANE AND EXTRACELLULAR MATRIX”. In:Mathematical biosciences and engineering : MBE 10.1 (Feb. 2013), pp. 75–101. ISSN: 1547-1063. doi:10.3934/mbe.2013.10.75. URL: https://www.ncbi.nlm.nih.gov/pmc/articles/PMC6557636/ (visited on 12/20/2024).

[57] Yafei Wang et al. “Impact of tumor-parenchyma biomechanics on liver metastatic progression: a multi-model approach”. In:Scientific Reports 11.1 (Jan. 18, 2021), p. 1710. ISSN: 2045-2322. doi:10.1038/s41598-020-78780-7. URL: https://www.nature.com/articles/s41598-020-78780-7 (visited on 12/20/2024).

[58] Jacob G. Scott et al. “Spatial Metrics of Tumour Vascular Organisation Predict Radiation Efficacy in a Computational Model”. In:PLOS Computational Biology 12.1 (Jan. 22, 2016). Publisher: Public Library of Science, e1004712. ISSN: 1553-7358. doi:10.1371/journal.pcbi.1004712. URL: https://journals.plos.org/ploscompbiol/article?id=10.1371/journal.pcbi.1004712 (visited on 12/02/2024).

[59] Rifat Atun et al. “Expanding global access to radiotherapy”. In:The Lancet Oncology 16.10 (Sept. 1, 2015). Publisher: Elsevier, pp. 1153–1186. ISSN: 1470-2045, 1474-5488. doi:10.1016/S1470-2045(15)00222-3. URL: https://www.thelancet.com/journals/lanonc/article/PIIS1470-2045(15)00222-3/fulltext (visited on 02/19/2025).

[60] David A. Jaffray et al. “Harnessing progress in radiotherapy for global cancer control”. In: Nature Cancer 4.9 (Sept. 2023). Publisher: Nature Publishing Group, pp. 1228–1238. ISSN: 2662-1347. doi:10.1038/s43018-023-00619-7. URL: https://www.nature.com/articles/s43018-023-00619-7 (visited on 02/19/2025).

[61] Gibin G. Powathil, Douglas J. A. Adamson, and Mark A. J. Chaplain. “Towards Predicting the Response of a Solid Tumour to Chemotherapy and Radiotherapy Treatments: Clinical Insights from a Computational Model”. In:PLOS Computational Biology 9.7 (July 11, 2013). Publisher: Public Library of Science, e1003120. ISSN: 1553-7358. doi:10.1371/journal.pcbi.1003120. URL: https://journals.plos.org/ploscompbiol/article?id=10.1371/journal.pcbi.1003120 (visited on 12/18/2024).

[62] Stephen Joseph McMahon. “The linear quadratic model: usage, interpretation and challenges”. In:Physics in Medicine & Biology 64.1 (Dec. 2018). Publisher: IOP Publishing, 01TR01. ISSN: 0031-9155. doi:10.1088/1361-6560/aaf26a. URL: https://dx.doi.org/10.1088/1361-6560/aaf26a (visited on 12/18/2024).

[63] Gibin Powathil et al. “Modeling the Spatial Distribution of Chronic Tumor Hypoxia: Implications for Experimental and Clinical Studies”. In:Computational and Mathematical Methods in Medicine 2012.1 (2012). eprint: https://onlinelibrary.wiley.com/doi/pdf/10.1155/2012/410602, p. 410602. ISSN: 1748-6718. doi:10.1155/2012/410602. URL: https://onlinelibrary.wiley.com/doi/abs/10.1155/2012/410602 (visited on 12/18/2024).

[64] Alexandru Daşu, Iuliana Toma-Daşu, and Mikael Karlsson. “The effects of hypoxia on the theoretical modelling of tumour control probability”. In:Acta Oncologica 44.6 (Jan. 1, 2005). Publisher: Taylor & Francis eprint: https://doi.org/10.1080/02841860500244435, pp. 563– 571. ISSN: 0284-186X. doi:10.1080/02841860500244435. URL: https://doi.org/10.1080/02841860500244435 (visited on 12/18/2024).

[65] Bradly G. Wouters and J. Martin Brown. “Cells at Intermediate Oxygen Levels Can Be More Important Than the “Hypoxic Fraction” in Determining Tumor Response to Fractionated Radiotherapy”. In:Radiation Research 147.5 (May 1, 1997), pp. 541–550. ISSN: 0033-7587. doi:10.2307/3579620. URL: https://doi.org/10.2307/3579620 (visited on 12/18/2024).

[66] Tikvah Alper and P. Howard-Flanders. “Role of Oxygen in Modifying the Radiosensitivity of E. Coli B.” In:Nature 178.4540 (Nov. 1956). Publisher: Nature Publishing Group, pp. 978– 979. ISSN: 1476-4687. doi:10.1038/178978a0. URL: https://www.nature.com/articles/178978a0 (visited on 12/18/2024).

[67] Juan Uriel Legaria-Peña, Félix Sánchez-Morales, and Yuriria Cortés-Poza. “Understanding post-angiogenic tumor growth: Insights from vascular network properties in cellular automata modeling”. In:Chaos, Solitons & Fractals186 (Sept. 1, 2024), p. 115199. ISSN: 0960-0779. doi:10.1016/j.chaos.2024.115199. URL: https://www.sciencedirect.com/science/article/pii/S0960077924007513 (visited on 12/17/2024).

[68] Joshua A. Bull et al. MuSpAn: A Toolbox for Multiscale Spatial Analysis. Pages: 2024.12.06.627195 Section: New Results. Dec. 8, 2024. doi:10.1101/2024.12.06.627195. URL: https://www.biorxiv.org/content/10.1101/2024.12.06.627195v1 (visited on 12/18/2024).

[69] Joshua A. Bull et al. “Extended correlation functions for spatial analysis of multiplex imaging data”. In:Biological Imaging4 (Jan. 2024), e2. ISSN: 2633-903X. doi:10.1017/S2633903X24000011. URL: https://www.cambridge.org/core/journals/biological-imaging/article/extended-correlation-functions-for-spatial-analysis-of-multiplex-imaging-data/FB677F0E100658E36725C5B4A3944EB7 (visited on 12/18/2024).

[70] Joshua A. Bull et al. Integrating diverse statistical methods to analyse stage-discriminatory cell interactions in colorectal neoplasia. Pages: 2024.06.02.597010 Section: New Results. June 3, 2024. doi:10.1101/2024.06.02.597010. URL: https://www.biorxiv.org/content/10.1101/2024.06.02.597010v1 (visited on 12/18/2024).

[71] John A. Fozard et al. “Techniques for analysing pattern formation in populations of stem cells and their progeny”. In:BMC Bioinformatics 12.1 (Oct. 12, 2011), p. 396. ISSN: 1471-2105. doi:10.1186/1471-2105-12-396. URL: https://doi.org/10.1186/1471-2105-12-396 (visited on 12/18/2024).

[72] S. Dini, B. J. Binder, and J. E. F. Green. “Understanding interactions between populations: Individual based modelling and quantification using pair correlation functions”. In:Journal of Theoretical Biology439 (Feb. 14, 2018), pp. 50–64. ISSN: 0022-5193. doi:10.1016/j.jtbi.2017.11.014. URL: https://www.sciencedirect.com/science/article/pii/S0022519317305210 (visited on 12/18/2024).

[73] Fergus R Cooper et al. “Chaste: Cancer, Heart and Soft Tissue Environment”. In: Journal of open source software 5.47 (Mar. 13, 2020), p. 1848. ISSN: 2475-9066. doi: 10.21105/joss.01848. URL: https://www.ncbi.nlm.nih.gov/pmc/articles/PMC7614534/ (visited on 12/18/2024).

[74] Gary R. Mirams et al. “Chaste: An Open Source C++ Library for Computational Physiology and Biology”. In: PLOS Computational Biology 9.3 (Mar. 14, 2013). Publisher: Public Library of Science, e1002970. ISSN: 1553-7358. doi: 10.1371/journal.pcbi.1002970. URL: https://journals.plos.org/ploscompbiol/article?id=10.1371/journal.pcbi.1002970 (visited on 12/18/2024).

[75] Joe Pitt-Francis et al. “Chaste: A test-driven approach to software development for biological modelling”. In: Computer Physics Communications. 40 YEARS OF CPC: A celebratory issue focused on quality software for high performance, grid and novel computing architectures 180.12 (Dec. 1, 2009), pp. 2452–2471. ISSN: 0010-4655. doi: 10.1016/j.cpc.2009.07.019. URL: https://www.sciencedirect.com/science/article/pii/S0010465509002604 (visited on 12/18/2024).

[76] Adrian Baddeley, Imre Bárány, and Rolf Schneider. “Spatial point processes and their applications”. In: Stochastic Geometry: Lectures Given at the CIME Summer School Held in Martina Franca, Italy, September 13–18, 2004 (2007). Publisher: Springer, pp. 1–75.

[77] H. Edelsbrunner, D. Kirkpatrick, and R. Seidel. “On the shape of a set of points in the plane”. In: IEEE Transactions on Information Theory 29.4 (July 1983). Conference Name: IEEE Transactions on Information Theory, pp. 551–559. ISSN: 1557-9654. doi: 10.1109/TIT.1983.1056714. URL: https://ieeexplore.ieee.org/abstract/document/1056714 (visited on 12/18/2024).

[78] Wentao Sui and Dan Zhang. “Four Methods for Roundness Evaluation”. In: Physics Procedia. International Conference on Applied Physics and Industrial Engineering 2012 24 (Jan. 1, 2012), pp. 2159–2164. ISSN: 1875-3892. doi: 10.1016/j.phpro.2012.02.317. URL: https://www.sciencedirect.com/science/article/pii/S1875389212003604 (visited on 12/18/2024).

[79] Robert M. Haralick. “A Measure for Circularity of Digital Figures”. In: IEEE Transactions on Systems, Man, and Cybernetics SMC- 4.4 (July 1974), pp. 394–396. ISSN: 2168-2909. doi: 10.1109/TSMC.1974.5408463. URL: https://ieeexplore.ieee.org/document/5408463/?arnumber=5408463 (visited on 12/13/2024).

[80] T. Alarćon, H. M. Byrne, and P. K. Maini. “A cellular automaton model for tumour growth in inhomogeneous environment”. In: Journal of Theoretical Biology 225.2 (Nov. 21, 2003), pp. 257–274. ISSN: 0022-5193. doi: 10.1016/S0022-5193(03)00244-3. URL: https://www.sciencedirect.com/science/article/pii/S0022519303002443 (visited on 12/17/2024).

[81] J. A. Grogan et al. “Simulating tumour vasculature at multiple scales”. In: Proceedings of the 2014 6th International Advanced Research Workshop on In Silico Oncology and Cancer Investigation - The CHIC Project Workshop (IARWISOCI). Proceedings of the 2014 6th International Advanced Research Workshop on In Silico Oncology and Cancer Investigation - The CHIC Project Workshop (IARWISOCI). Nov. 2014, pp. 1–4. doi: 10.1109/IARWISOCI.2014.7034633. URL: https://ieeexplore.ieee.org/abstract/document/7034633 (visited on 12/18/2024).

[82] James A. Grogan et al. “Microvessel Chaste: An Open Library for Spatial Modeling of Vascularized Tissues”. In: Biophysical Journal 112.9 (May 9, 2017). Publisher: Elsevier, pp. 1767– 1772. ISSN: 0006-3495, 1542-0086. doi: 10.1016/j.bpj.2017.03.036. URL: https://www.cell.com/biophysj/abstract/S0006-3495(17)30384-3 (visited on 12/18/2024).

[83] Boer Deng et al. “Biological role of matrix stiffness in tumor growth and treatment”. In: Journal of Translational Medicine 20.1 (Nov. 22, 2022), p. 540. ISSN: 1479-5876. doi: 10.1186/s12967-022-03768-y. URL: https://doi.org/10.1186/s12967-022-03768-y (visited on 02/19/2025).

[84] Ludivine Guillaume et al. “Characterization of the physical properties of tumor-derived spheroids reveals critical insights for pre-clinical studies”. In: Scientific Reports 9.1 (Apr. 29, 2019). Publisher: Nature Publishing Group, p. 6597. ISSN: 2045-2322. doi: 10.1038/s41598-019-43090-0. URL: https://www.nature.com/articles/s41598-019-43090-0 (visited on 02/19/2025).

[85] Pauline Chassonnery et al. “Fibre crosslinking drives the emergence of order in a three-dimensional dynamical network model”. In: Royal Society Open Science 11.1 (Jan. 31, 2024), p. 231456. doi: 10.1098/rsos.231456. URL: https://royalsocietypublishing.org/doi/10.1098/rsos.231456 (visited on 12/25/2024).

[86] Juan Arellano-Tintó et al. Multiscale modelling shows how cell-ECM interactions impact ECM fibre alignment and cell detachment. Dec. 11, 2024. doi: 10.1101/2024.12.05.627121. URL: https://www.biorxiv.org/content/10.1101/2024.12.05.627121v2 (visited on 12/25/2024).

[87] James W. Reinhardt and Keith J. Gooch. “An Agent-Based Discrete Collagen Fiber Network Model of Dynamic Traction Force-Induced Remodeling”. In: Journal of Biomechanical Engineering 140.51003 (Feb. 22, 2018). ISSN: 0148-0731. doi: 10.1115/1.4037947. URL: https://doi.org/10.1115/1.4037947 (visited on 12/25/2024).

[88] Yu Zheng et al. “Modeling cell migration regulated by cell extracellular-matrix micromechanical coupling”. In: Physical Review E 100.4 (Oct. 11, 2019), p. 043303. ISSN: 2470-0045, 2470-0053. doi: 10.1103/PhysRevE.100.043303. URL: https://link.aps.org/doi/10.1103/PhysRevE.100.043303 (visited on 12/25/2024).

[89] Rebecca M. Crossley et al. “Modeling the extracellular matrix in cell migration and morphogenesis: a guide for the curious biologist”. In: Frontiers in Cell and Developmental Biology 12 (Mar. 1, 2024). Publisher: Frontiers. ISSN: 2296-634X. doi: 10.3389/fcell.2024.1354132. URL: https://www.frontiersin.org/journals/cell-and-developmental-biology/articles/10.3389/fcell.2024.1354132/full (visited on 12/18/2024).

[90] Heber L. Rocha et al. “A multiscale model of immune surveillance in micrometastases gives insights on cancer patient digital twins”. In: npj Systems Biology and Applications 10.1 (Dec. 4, 2024), pp. 1–12. ISSN: 2056-7189. doi: 10.1038/s41540-024-00472-z. URL: https://www.nature.com/articles/s41540-024-00472-z (visited on 12/20/2024).

[91] Mark W. Dewhirst and Timothy W. Secomb. “Transport of drugs from blood vessels to tumour tissue”. In: Nature Reviews Cancer 17.12 (Dec. 2017). Publisher: Nature Publishing Group, pp. 738–750. ISSN: 1474-1768. doi: 10.1038/nrc.2017.93. URL: https://www.nature.com/articles/nrc.2017.93 (visited on 12/18/2024).

[92] Ruth Ganss. “Tumour vessel remodelling: new opportunities in cancer treatment”. In: (Feb. 6, 2020). Section: Vascular Biology. doi: 10.1530/VB-19-0032. URL: https://vb.bioscientifica.com/view/journals/vb/2/1/VB-19-0032.xml (visited on 12/18/2024).

[93] Jie Ma and David J. Waxman. “Combination of antiangiogenesis with chemotherapy for more effective cancer treatment”. In: Molecular Cancer Therapeutics 7.12 (Dec. 15, 2008), pp. 3670– 3684. ISSN: 1535-7163. doi: 10.1158/1535-7163.MCT-08-0715. URL: https://doi.org/10.1158/1535-7163.MCT-08-0715 (visited on 12/18/2024).

[94] Sabrina M. Lewis et al. “Spatial omics and multiplexed imaging to explore cancer biology”. In: Nature Methods 18.9 (Sept. 2021). Publisher: Nature Publishing Group, pp. 997–1012. ISSN: 1548-7105. doi: 10.1038/s41592-021-01203-6. URL: https://www.nature.com/articles/s41592-021-01203-6 (visited on 12/13/2024).

[95] David Robert Grimes et al. “A method for estimating the oxygen consumption rate in multicellular tumour spheroids”. In: Journal of The Royal Society Interface 11.92 (Mar. 6, 2014). Publisher: Royal Society, p. 20131124. doi:10.1098/rsif.2013.1124. URL: https://royalsocietypublishing.org/doi/full/10.1098/rsif.2013.1124 (visited on 12/18/2024).

[96] Sophie Laget et al. “Technical Insights into Highly Sensitive Isolation and Molecular Characterization of Fixed and Live Circulating Tumor Cells for Early Detection of Tumor Invasion”. In: PLOS ONE 12.1 (Jan. 6, 2017). Publisher: Public Library of Science, e0169427. ISSN: 1932-6203. doi: 10.1371/journal.pone.0169427. URL: https://journals.plos.org/plosone/article?id=10.1371/journal.pone.0169427 (visited on 12/18/2024).

[97] P. Pathmanathan et al. “A computational study of discrete mechanical tissue models”. In: Physical Biology 6.3 (Apr. 2009), p. 036001. ISSN: 1478-3975. doi: 10.1088/1478-3975/6/3/036001. URL: https://dx.doi.org/10.1088/1478-3975/6/3/036001 (visited on 12/18/2024).

